# A next generation of hierarchical Bayesian analyses of hybrid zones enables direct quantification of variation in introgression in R

**DOI:** 10.1101/2024.03.29.587395

**Authors:** Zachariah Gompert, Devon A. DeRaad, C. Alex Buerkle

## Abstract

Hybrid zones, where genetically distinct groups of organisms meet and interbreed, offer valuable insights into the nature of species and speciation. Here, we present a new R package bgchm, for population genomic analyses of hybrid zones. This R package extends and updates the existing bgc software and combines Bayesian analyses of hierarchical genomic clines with Bayesian methods for estimating hybrid indexes, interpopulation ancestry proportions, and geographic clines. Compared to existing software, bgchm offers enhanced efficiency through Hamiltonian Monte Carlo sampling and the ability to work with genotype likelihoods combined with a hierarchical Bayesian approach, enabling accurate inference for diverse types of genetic datasets. The package also facilitates the quantification of introgression patterns across genomes, which is crucial for understanding reproductive isolation and speciation genetics. We first describe the models underlying bgchm and then provide an overview of the R package and illustrate its use through the analysis of simulated and empirical data sets. We show that bgchm generates accurate estimates of model parameters under a variety of conditions, especially when the genetic loci analyzed are highly ancestry informative. This includes relatively robust estimates of genome-wide variability in clines, which has not been the focus of previous models and methods. We also illustrate how both selection and genetic drift contribute to variability in introgression among loci and how additional information can be used to help distinguish these contributions. We conclude by describing the promises and limitations of bgchm, comparing bgchm to other software for genomic cline analyses, and identifying areas for fruitful future development.

## Introduction

Hybrid zones form when genetically distinct groups of organisms meet, mate and produce offspring (Barton & Hewitt, 1985; Gompert & Buerkle, 2016). Studies of hybrid zones provide powerful opportunities to analyze interactions between divergent gene pools in the wild (Barton *et al*., 1993; Buerkle & Lexer, 2008; Gompert *et al*., 2017), and are especially relevant for testing hypotheses about the nature and genetic basis of species and speciation (Harrison & Larson, 2014; Firneno *et al*., 2023). The ease with which genomic data can be generated has vastly increased the potential for genomic analyses of hybrid zones. Simultaneously, advances in analytical approaches and computer software packages have increased the ability of investigators to make evolutionary inferences from hybrid zone data (reviewed in Gompert *et al*., 2017).

Hybrid zone theory was largely developed in the mid to late 1900s (e.g., Haldane, 1948; Endler, 1977; Barton, 1979, 1983; Barton & Hewitt, 1985). Results from this body of theory provide a means to connect model parameters describing the width, location and shape of geographic clines in hybrid zones to evolutionary parameters and processes, such as selection and dispersal (Barton & Hewitt, 1985). Such geographic cline approaches have been used extensively and productively in speciation research (e.g., Szymura & Barton, 1986; Mallet *et al*., 1990; Dasmahapatra *et al*., 2002; Carling & Brumfield, 2008; Teeter *et al*., 2008; Westram *et al*., 2021; Caeiro-Dias *et al*., 2023). Nonetheless, these approaches are not always applicable, especially when hybridization lacks a major geographic axis (e.g., Harrison & Rand, 1989; Rieseberg *et al*., 1999; Mandeville *et al*., 2015; Chaturvedi *et al*., 2020), and are but one of several windows into the evolutionary processes provided by hybrid zones.

The prevalence and genomic composition of hybrids in hybrid zones provides additional information about the strength and form of reproductive isolation (Jiggins & Mallet, 2000). Moreover, genomic approaches can go beyond simple classification of hybrids as F1s, F2s, or backcrosses by describing hybrid genomes quantitatively. For example, genome composition can be measured with a hybrid index, which denotes the proportion of an individual’s genome inherited from one of two hybridizing lineages (Buerkle, 2005) (akin to the admixture proportions from structure, Pritchard *et al*., 2000), and by an interpopulation (i.e., interclass) ancestry proportion, which indicates the proportion of an individual’s genome with gene copies inherited from both hybridizing species (Gompert & Buerkle, 2010; Fitzpatrick, 2012; Gompert *et al*., 2014; Shastry *et al*., 2021). Together these metrics provide flexible, continuous summaries of the genetic makeup of hybrids that are relevant for understanding hybrid zone dynamics (e.g., interpopulation ancestry will be high when matings between non-admixed individuals or between hybrids and non-admixed individuals are common).

Finally, genomic cline models can be used to quantify introgression from one genomic background to another, with a focus on patterns of heterogeneity in introgression across the genome and associated evolutionary processes (Szymura & Barton, 1986; Gompert & Buerkle, 2011; Fitzpatrick, 2013b). In this context, recombination and independent assortment in hybrids create new genotypic combinations that are subject to selection based on their effects on hybrid fitness. Such selection, along with other factors (e.g., patterns of linkage disequilibrium) and processes (e.g., recombination, drift, gene flow, etc.), affect patterns of introgression in hybrid zones (Barton, 1983; Gompert *et al*., 2012b; Lindtke & Buerkle, 2015; Schumer *et al*., 2018; McFarlane *et al*., 2021). Consequently, outcomes of hybridization and patterns of introgression often vary across the genome (e.g., Nolte *et al*., 2009; Larson *et al*., 2013; Sung *et al*., 2018; Chaturvedi *et al*., 2020; Wagner *et al*., 2020; Caeiro-Dias *et al*., 2023), which can provide additional information about the genetics of speciation (Payseur, 2010; Harrison & Larson, 2016; Gompert *et al*., 2017). This variation can be quantified using genomic cline models and compared across sets of loci, chromosomes, and hybrid zones with implications for understanding the genetics of reproductive isolation, the repeatability of speciation, and coupling of barrier loci in hybrid zones (e.g., Teeter *et al*., 2010; Larson *et al*., 2013; Taylor *et al*., 2014; Nikolakis *et al*., 2022; Firneno *et al*., 2023; McFarlane *et al*., 2023).

Here, we present a new R package bgchm, which combines Bayesian analyses of hierarchical genomic clines with Bayesian methods for (i) estimating hybrid indexes and interpopulation ancestry proportions and (ii) fitting geographic cline models. This package builds on the foundation of the existing bgc software (Gompert & Buerkle, 2012), but replaces the Barton cline model with the logit-logistic cline model proposed by Fitzpatrick (2013b). We describe the details of the models and software usage below, but here briefly highlight some of the most salient aspects of this R package (we make detailed comparisons with related software in the Discussion). First, bgchm replaces traditional Metropolis-Hastings Markov chain Monte Carlo with Hamiltonian Monte Carlo, which generally results in much more efficient sampling of posterior distributions (Neal *et al*., 2011). As with the original bgc, bgchm retains the ability to analyze data comprising known genotypes or to work directly with genotype likelihoods, which are standard output of most modern genetic variant callers and imputation methods. This makes it possible to account for uncertainty in genotypes in analyses and is critical for accurate and powerful inference from low to moderate coverage DNA sequence data sets. bgchm additionally adds the option to work directly with local (locus-specific) ancestry estimates instead of genotypic data. Finally, bgchm retains a hierarchical Bayesian approach to cline inference. Together, these features result in much more reliable inference of cline standard deviation parameters, which provide high-level summaries of introgression across the genome and are relevant for studying coupling in hybrid zones (Firneno *et al*., 2023). Additionally, by separately estimating hybrid indexes and clines, but still retaining the hierarchical structure of the model, bgchm drastically improves parallelization relative to bgc and also allows comparisons among different sets of loci (e.g., trait associated versus putatively neutral loci, or different chromosomes) without assuming all loci in a set share the same cline parameters. These features are important for scaling cline analyses to genome-level data sets.

In this manuscript, we first describe the core models underlying bgchm. We then provide an overview of the R package usage and illustrate its use through the analysis of simulated data sets. Where relevant, we compare bgchm to HIest, which provided the original implementation of the logit-logistic cline model. We further demonstrate the usage of bgchm via the analysis of a butterfly hybrid zone data set. We conclude by discussing the potential and limitations of bgchm, comparing this R package with other hybrid zone analysis software, and identifying possibilities for further developments.

## Methods

### Models

We consider three sets of models to describe genomic patterns of admixture and introgression in hybrid zones, specifically, models to infer hybrid indexes, ancestry class proportions, and genomic clines (geographic clines models are described in the the Online Supplemental Materials [OSM]). We first describe these models for the case where genotypes are assumed to be known without error before presenting extensions for modeling genotype uncertainty or working directly with local ancestry estimates.

#### Hybrid index model

We follow the basic structure of the hybrid index model proposed by (Buerkle, 2005). Here, hybrid indexes are defined with respect to two putative source or reference populations defined to represent or approximate the genetic composition of two hybridizing species or lineages. The hybrid index for individual *j*, *H_j_*, denotes the proportion of individual *j*’s genome that is best modeled as being inherited from one of the two source populations (labeled source 1). Consequently, 1 *− H_j_* denotes the proportion of the genome inherited from the other source population (labeled source 0). Hybrid indexes are based on supervised learning of allele frequencies within source populations that are defined *a priori* and are equivalent to admixture proportions estimated in an unsupervised learning context (Pritchard *et al*., 2000; Gompert *et al*., 2014). Here we consider only two source populations. We assume that the genotypic data for individual *j* and locus *i* is binomially distributed conditional on ancestry of the alleles at locus *i* and corresponding parental allele frequencies (*P*_0*i*_ and *P*_1*i*_), and similarly that the ancestry at locus *i* is binomially distributed conditional on the hybrid index, *H_j_*. This results in the following likelihood model for estimating hybrid indexes:

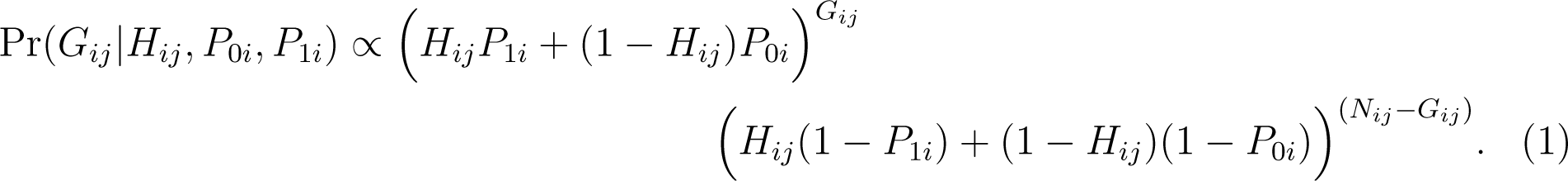

Here, *G_ij_*is the count of one of two alleles (e.g., the non-reference allele) and *N_ij_* denotes the number of allele copies for the individual and locus (i.e., two for diploids). We assume a beta prior on hybrid indexes, such that *H_j_∼* beta(*α* = 0.5*, β* = 0.5), which corresponds with Jeffery’s minimally informative prior.

#### Ancestry class proportions model

Our model for ancestry class proportions is similar to the interpopulation ancestry models described by Gompert *et al*. (2014) and Shastry *et al*. (2021) (i.e., the Q model). However, unlike these models, our ancestry class proportions model assumes source populations are defined with known allele frequencies *a priori* (i.e., supervised learning, as in our hybrid index model). We designate the ancestry class proportions *Q*_00_, *Q*_11_ and *Q*_10_ to denote (i) the proportion of an individual’s genome where both gene copies were inherited from source (i.e., reference) population 0 (*Q*_00_), (ii) the proportion of an individual’s genome where both gene copies were inherited from source population 1 (*Q*_11_), (iii) the proportion of an individual’s genome where one gene copy was inherited from each source population (*Q*_10_, i.e., interpopulation ancestry). The main purpose of the model is to estimate these ancestry class proportions. Note that hybrid index can be derived directly from the ancestry class proportions as 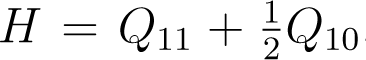. We define a likelihood analogous to Eqn. 1 for the ancestry class proportions as:

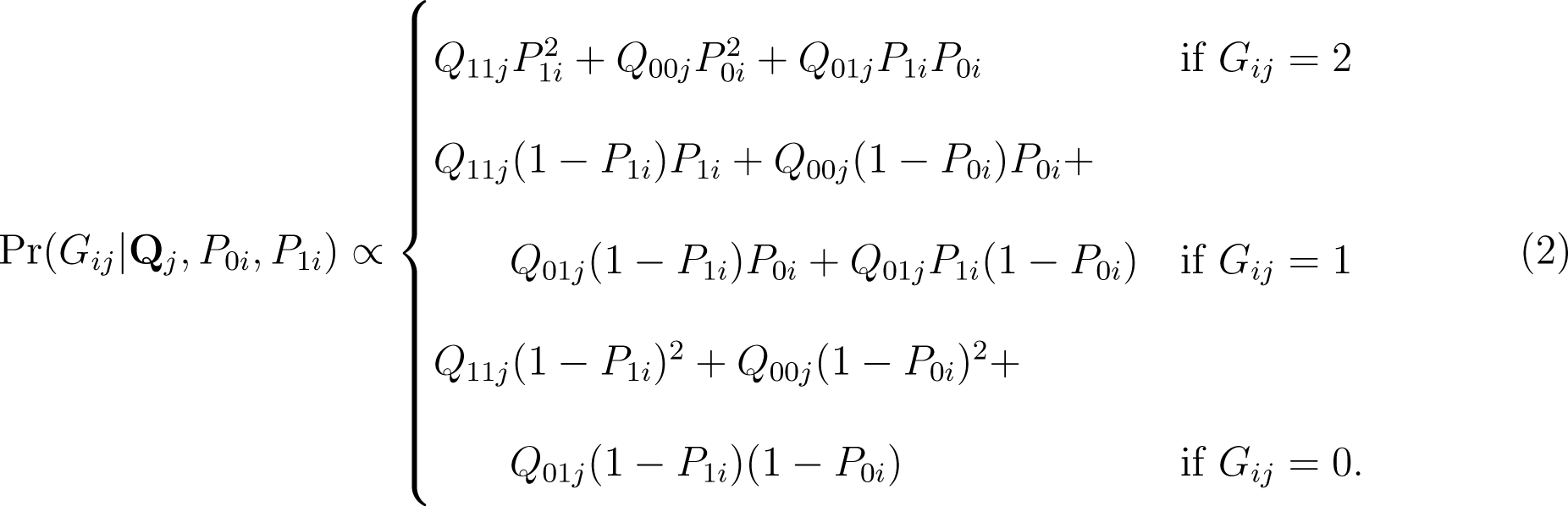

#### Genomic clines model

Genomic clines represent the probability of locus-specific ancestry along a genomewide admixture gradient, that is, as a function of hybrid index (Szymura & Barton, 1986; Gompert & Buerkle, 2009, 2011). Here, we model genomic clines with the logit-logistic model proposed by Fitzpatrick (2013b). With this function, the probability that a gene copy for locus *i* and individual *j* was inherited from source population 1 (as opposed to source population 0) is 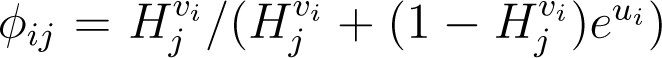, where *H_j_* is the hybrid index (i.e., the proportion of the genome inherited from population 1), *v_i_* gives the slope of the cline for locus *i* relative to the genome-average (*v̄* = 1) and *u_i_* specifies the center of the cline for locus *i* relative to both the genome average and *v_i_* (Fitzpatrick, 2013b). We use the re-parameterization from Bailey (2022) and Firneno *et al*. (2023) that defines 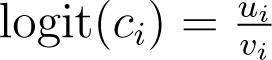 to specify a more intuitive cline center parameter (*c_i_*), which indicates the hybrid index value at which *ϕ_ij_* = 0.5, that is where the probability of inheriting an allele from each source population is equal. Genomic cline slopes greater than 1 indicate a steeper cline than the admixture gradient, whereas clines less than 1 indicate a shallower cline. Similarly, centers greater than 0.5 indicate an overall excess of source 0 ancestry, whereas centers less than 0.5 indicate an excess of source 1 ancestry.

We specify the following likelihood model for the genetic data at locus *i* specified in terms of *ϕ_ij_*, which is itself a function of hybrid index (*H_j_*, a property of an individual) and the cline parameters *v_i_* and *c_i_* (properties of a locus):

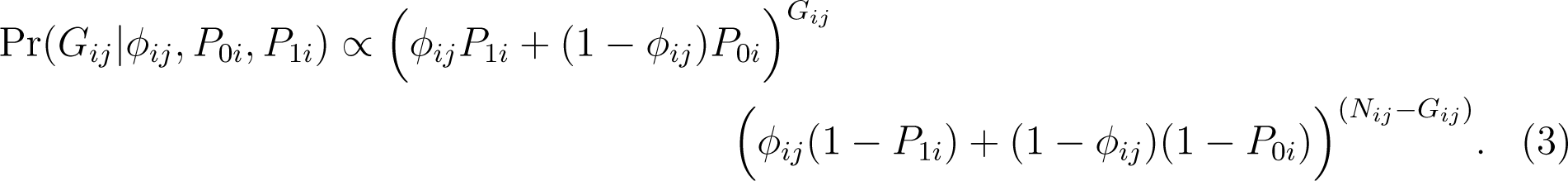

Note the similarity between Eqn. 3 and Eqn. 1; the forms are identical except that *ϕ_ij_* (the probability of ancestry from source 1 for locus *i* and individual *j*) in Eqn. 3 replaces *H_j_* (the marginal probability of ancestry from source 1 for individual *j*) in Eqn. 1.

Following Gompert & Buerkle (2011) and Firneno *et al*. (2023), we define hierarchical priors for the cline parameters *v_i_* and *c_i_*. Hierarchical modelling allows information on the genomic variability of introgression to be shared across loci and explicitly acknowledges the partial dependence (and partial independence) of introgression across the genome (Gompert & Buerkle, 2011; Betancourt & Girolami, 2015). In general, hierarchical modelling in such cases is conceptually preferable to the alternative assumptions of complete independence of units (e.g., introgression patterns across loci) as implied by fixed, independent priors, or the complete lack of independence among units as implied by a shared parameter for all units (e.g., the same cline parameters for all loci; Gelman *et al*., 1995; Fordyce *et al*., 2011). In our standard model, we specify the following priors for cline parameters (modifications are discussed below): log_10_(*v_i_*) *∼* normal(*µ* = 0*, σ* = *σ_v_*) and logit(*c_i_*) *∼* normal(*µ* = 0*, σ* = *σ_c_*). The log and logit functions are used to set the expected means of *v_i_*and *c_i_* to 0 (log_10_(1) = 0 and logit(0.5) = 0) and also to project these parameters onto the scale of *−∞* and *∞*. The means of both priors are set to 0 to reflect the fact that, assuming the same loci (or random subsets of the same loci) are used to infer hybrid indexes and to fit genomic clines, the average deviation of locus-specific clines from the genome-average should by definition be 0 (Gompert & Buerkle, 2011). This can be relaxed in cases where distinct sets of loci are used for cline fitting and hybrid indexes, as we discuss below. Such a zero-centered prior does not enforce a hard sum-to-zero constraint, but rather serves as a form of soft centering. We discuss hard-centering (i.e., sum-to-zero constraints) in the ‘Software usage” section.

The standard deviation parameters, *σ_v_* and *σ_c_*describe the variability of cline slopes and centers across the genome, and can be related to the extent of coupling among loci (Barton, 1983; Firneno *et al*., 2023). These standard deviation parameters simultaneously inform and are informed by the locus-specific cline parameters and it is this co-dependency that allows information sharing across loci. As such, these cline standard deviations (*σ_v_* and *σ_c_*) are estimated from the data as part of the analysis (at least in the standard model, modifications to this procedure are discussed below). Thus, priors (hyperpriors) are placed on the standard deviations, *σ_v_ ∼* normal(*µ* = 0*, σ* = *σ*_0_) and *σ_c_ ∼* normal(*µ* = 0*, σ* = *σ*_0_), with *σ*_0_ set by users.

#### Alternative model specifications and assumptions

Having described a standard version of each of our models for hybrid indexes, ancestry class proportions and genomic clines above, we now discuss modifications and variants of these models. First, the model descriptions above assume that genotypes are known without error. However, modern sequencing technologies and bioinformatic tools generate finite numbers of reads or sequences covering each segment of DNA, uncertain base calls and mapping errors. These sources of uncertainty mean that genotypes are often uncertain. This is reflected in the genotype likelihoods output by most variant calling software (e.g., samtools and bcftools; Li, 2011). Uncertainty can also arise from genotype imputation or Bayesian inference of genotypes (in these cases uncertainty is often captured in a posterior probability rather than a likelihood, but can be incorporated in the same manner). Thus, we include modifications of all three core models to incorporate uncertainty in genotypes by working directly with relative likelihoods or posterior probabilities of genotypes (e.g., as output by some Bayesian genotype inference methods, e.g., Shastry *et al*., 2021). In such cases, the likelihoods given in Eqns. 1–3 are replaced by the average likelihood of the parameter values conditional on each genotype and weighted by relative genotype likelihoods or posterior probabilities. For example, Eqn. 3 becomes Pr(*G_ij_|ϕ_ij_, P*_0_*_i_, P*_1_*_i_*) = Pr(*G_ij_* = 0*|ϕ_ij_, P*_0_*_i_, P*_1_*_i_*)Pr(*G_ij_* = 0) + Pr(*G_ij_* = 1*|ϕ_ij_, P*_0_*_i_, P*_1_*_i_*)Pr(*G_ij_* = 1) + Pr(*G_ij_* = 2*|ϕ_ij_, P*_0_*_i_, P*_1_*_i_*)Pr(*G_ij_* = 2).

We also consider alternative models where we assume locus-specific ancestry is itself known or has been estimated using one of many programs designed for local-ancestry inference (e.g., Li & Stephens, 2003; Maples *et al*., 2013; Browning *et al*., 2023). With known local ancestry, the likelihood equations no longer depend on parental allele frequencies, and, for example, Eqn. 3 can be simplified to:

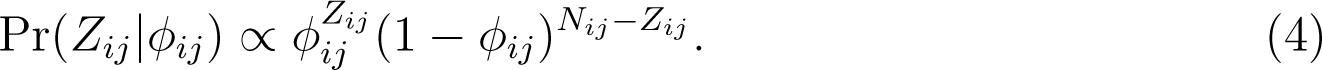

Here, *Z_ij_* denotes the ancestry of locus *i* in individual *j*, that is the number of gene copies (out of *N_ij_*) individual *j* inherited from source population 1 at this specific locus.

We define two additional variants of the genomic clines model, both of which can be applied with known genotypes, uncertain genotypes, or local ancestry. First, one such variant allows the standard deviations of the hierarchical priors, *σ_v_* and *σ_c_*, to be specified and fixed. As described in more detail in the “Software usage” section, this makes it possible to first estimate these parameters using the standard hierarchical model based on a random subset of data, and then to fix these parameters for estimating clines for the full set of data enabling massive parallelization of the model fitting procedure across genetic loci. Alternatively, this model formulation can be used to specify weakly informative priors (i.e., relatively flat priors) and thereby implement a non-hierarchical version of the genomic clines model akin to Bailey (2022).

Second, in some cases, it can be useful to estimate the means of the hierarchical priors for *v* and *c* from the data rather than fix them at 0. As we illustrate in an example analysis below, this could be done if one estimates hybrid indexes based on one subset of loci (e.g., putative neutral regions of the genome, autosomes only, etc.) and then wants to ask whether a different subset of loci (e.g., trait-associated loci, other candidate genes, sex-linked loci, etc.) exhibit patterns of introgression that deviate on average from the subset of loci used for hybrid index inference. Consequently, we have also included models with unknown means for the hierarchical priors, log_10_(*v_i_*) *∼* normal(*µ* = *µ_v_, σ* = *σ_v_*) and logit(*c_i_*) *∼* normal(*µ* = *µ_c_, σ* = *σ_c_*). In such cases, we place normal priors on the unknown means as well as the unknown standard deviation, with *µ_v_ ∼* normal(*µ* = 0*, σ* = *µ*_0_) and *µ_c_ ∼* normal(*µ* = 0*, σ* = *µ*_0_), and with the prior standard deviation for these means, *µ*_0_, specified by the user.

### Software usage

We implemented the models described above in a new R package, bgchm, which updates the original bgc program (Gompert & Buerkle, 2012). The R package is available for direct installation from GitHub at https://github.com/zgompert/bgc-hm. The R package uses Stan (via rstan) for sampling from posteriors (Stan Development Team, 2022, 2024). This implementation makes it possible to fit the models using Hamiltonian Monte Carlo (HMC) rather than using more traditional Markov chain Monte Carlo algorithms. This is important as HMC routinely outperforms other algorithms especially in terms of more effectively exploring complex posterior distributions (Neal *et al*., 2011; Betancourt & Girolami, 2015). This means that far fewer HMC steps are generally required to obtain a good approximation of the posterior distribution and that the HMC algorithm is less likely to to stuck in one region of the posterior, especially when fitting hierarchical models and estimating higher-level standard deviations (Betancourt & Girolami, 2015). We specifically use the No-U-Turn Sampler (NUTS) from Stan (Hoffman *et al*., 2014; Betancourt, 2017). Integration with Stan and rstan also provides built-in diagnostics of HMC performance, including automated and standard warning message about performance and estimates of effective samples sizes and convergence diagnostics for each of the model parameters. Moreover, by using Stan, all of the HMC sampling is done based on compiled C++ code, rather than native R code, which is critical for minimizing the time required for model fitting.

The R package bgchm includes core functions for estimating hybrid indexes (est hi), ancestry class proportions (est Q), and genomic clines (est gencline). Each function is documented in the R package. The arguments to these functions determine which version of each model to fit, with each version corresponding to an internal compiled C++ program. Separate helper functions exist for estimating parental allele frequencies (est p) (this can also be done within the three core functions), summarizing posterior distributions (pp plot), and visualizing results (e.g., producing triangle interpopulation ancestry plots or plotting genomic clines). Additional functions for hierarchical geographic cline analyses are described in the OSM (these are not the main focus of the software, but are included for the convenience of users interested in hierarchical geographic cline models).

We have made the core functions modular for flexibility and ease of scaling, though this comes at the cost of not fully propagating uncertainty in parental allele frequencies and hybrid indexes in genomic cline analyses (this is an unfortunate but somewhat necessary trade-off). Consequently, the parental allele frequencies required for hybrid index, ancestry class proportions and genomic clines can be estimated within bgchm or provided from other software. Likewise, the hybrid indexes used in the genomic cline analysis can be estimated for all or a subset of loci and can be inferred withing bgchm or using other software (e.g., these could be admixture proportions from a model with *k* = 2 in structure, Pritchard *et al*., 2000). This set-up allows for extensive parallelization of cline fitting, and thus makes it possible to run bgchm on genome-scale data sets (assuming one has access to sufficient computational resources). Specifically, a standard analysis can begin by estimating hybrid indexes based on a moderate number of loci, several hundred to a thousand will generally be sufficient to obtain precise estimates of hybrid indexes. Then, for modest sized data sets (up to a few thousand loci and a few hundred hybrids), the full set of loci can be analyzed in a single hierarchical model (the standard model described above). For larger data sets (more than a thousand individuals or loci, and up to millions of SNPs), a subset of a few thousand loci can be fit in an initial hierarchical model to estimate the cline standard deviation parameters, *σ_v_* and *σ_c_*. These can then be fixed at their point estimates and the clines for the remaining loci can be fit in batches (and thus in parallel across CPUs or computer nodes) using these estimated standard deviations. This gains most of the benefit of using a hierarchical modelling framework without the cost of needing to fit clines for a very large number of loci in a single model. Additional parallelization is possible for all analyses by running multiple HMC chains in parallel (this is done within the bgchm program). We provide an example of batch parallelization of loci in the bgchm repository (https://github. com/zgompert/bgc-hm), including a UNIX shell script to control the batch parallelization.

As noted above in the “Model” section, the hierarchical prior structure for the standard genomic cline model results in soft centering of the cline parameters, such that the mean of the cline parameters (on the appropriate log or logit scale) is shrunk toward zero. However, this is not the same as a hard, sum-to-zero constraint, as implemented in the original bgc program (Gompert & Buerkle, 2012), which forces the mean of the cline parameters to be zero. We found that trying to enforce a hard sum-to-zero constraint within the HMC algorithm dramatically degraded performance of the algorithm. Moreover, a hard sum-to-zero constraint would only be possible when fitting all loci in a single model. We have thus instead opted to use soft centering, while also providing a function, sum2zero, that applies a sum-to-zero constraint to a set of cline parameter estimates after model fitting. This can be done based on the full HMC output (preferable when practical) or simply as an adjustment to the parameter estimates (useful when saving the full HCM output for all loci is computationally burdensome). Either of these options can be applied after batch processing of cline estimation for many loci, and thus makes it possible to apply a sum-to-zero constraint to genome-scale data. We think this is advisable in most cases, at least when the loci used to estimate hybrid indexes are the same or a random subset of those used to estimate clines. We think this because the genomic clines are, by definition, deviation from average introgression. In some cases, the soft centering might be sufficient to effectively constrain the mean of cline parameters to zero such that applying the hard sum-to-zero constraint is not necessary, but in other cases, including in the example empirical analysis we present, this is not true (this will depend—in ways that have yet to be fully investigated—on the distribution of hybrid indexes, variability of clines among loci, and other aspects of the data set and posterior distribution).

### Analyses of simulated data sets

We analyzed a series of simulated data sets to illustrate and evaluate the performance of the bgchm package. Aspects of this or related models have been analyzed extensively elsewhere and thus are not treated in depth here. For example, Gompert *et al*. (2012b) evaluated the concordance between loci with exceptional genomic cline parameters (from the original bgc model) and loci causally affecting fitness (this varies depending on the genetic architecture of fitness variation). Firneno *et al*. (2023) used simulations to assess the relationship between Barton’s theoretical coupling coefficient (*θ*) and the cline standard deviations from the hierarchical Bayesian logit-logistic model described here. Firneno *et al*. (2023) then quantified these cline standard deviations across a series of empirical data sets. Bailey (2022) examined the sensitivity of a non-hierarchical implementation of the logit-logistic cline model to the distribution of hybrid indexes. Here, our main focus is on demonstrating the general performance of the software and exploring specific aspects of these models or performance that have received less attention, including the effects of hierarchical modelling, allele frequency differences between parents, and our ability to accurately estimate cline standard deviations.

#### Analyses of simulations of hybrid indexes and ancestry class proportions

We first conducted simulations to evaluate the ability of bgchm to accurately estimate hybrid indexes and ancestry class coefficient from genetic data. Simulations were conducted using dfuse under a model of neutral secondary contact (Lindtke & Buerkle, 2015). The program dfuse implements individual-based simulations to model a hybrid zone that forms following secondary contact. The program tracks hybrid indexes, ancestry class proportions (specifically our *Q*_10_) and ancestry junctions along chromosomes. As such, it provides a way to simulate hybrids where the core parameters for these models, *H* and *Q*, are known. We conducted 50 replicate simulations of 200 generations where hybridization occurs in a single admixed deme with an adult carrying capacity of 500. The migration rate from the parental populations to the deme was set to 0.1. We simulated hermaphroditic, diploid organisms with ten chromosomes, each one Morgan in length. We output ancestry information for 51 loci spaced evenly along each of the ten chromosomes (510 loci total). At the end of each simulation, we randomly sampled 50 individuals from the hybrid zone deme for analysis. We then generated three genotypic data sets based on the output from each replicate simulation. Specifically, we sampled genotypes for each individual and locus based on the individual’s local ancestry and assumed parental allele frequencies of (i) 0 and 1, (ii) 0.25 and 0.75, (iii) or 0.45 an 0.55 for parents 0 and 1, respectively. This corresponds with parental allele frequency differences of 1 (fixed differences), 0.5 and 0.1. Genotypes were generated using binomial sampling (in R). From each of these genotypic data sets, we created an additional data set where the genotypes were uncertain. For this, we assumed the number of sequence reads for each individual and locus followed a Poisson distribution with *λ* = 7, and that sequences had a 1% error rate. Reads were sampled in R based on the genotypes and these parameters, and the likelihood of each genotype was then computed from the reads assuming the 1% error rate. Thus, for each of the 50 initial simulated hybrid zones, we generated six genetic data sets: parental allele frequency differences of 1, 0.5, or 0.1 with genotypes known or uncertain.

We then estimated hybrid indexes and ancestry class proportions with bgchm using the est hi and est Q functions. We did this using model for known genotypes or genotype likelihoods as appropriate, and with default HMC conditions for these functions: four HMC chains with 2000 steps, including 1000 warmup iterations. We used the known parental allele frequencies for the analysis. We summarized the posterior estimates of hybrid index and ancestry class proportions for each individual and simulated data set based on the posterior median (point estimate) and 90% credible intervals (CIs, specifically the 90% equal-tail probability intervals). We then evaluated performance by computing the mean absolute error (MAE) and the proportion of 90% CIs containing the true parameter value (90% CI coverage) for each data set.

#### Genomic cline analyses of simulated hybrid zones

We next conducted a series of simulations and analyses to evaluate the performance of the genomic cline models in bgchm. The first two sets of simulations were designed to evaluate the conditions under which bgchm could accurately estimate genomic cline parameters. Unlike hybrid indexes and ancestry class proportions, individual-based simulations, such as those in dfuse, do not generate known cline parameters. Thus, for these sets of simulations, we instead simulated hybrids using the logit-logistic genomic cline model as a generative model. The first set of simulations was designed to evaluate the effect of cline variability, that is variability in introgression across the genome, on our ability to accurately estimate cline parameters. For this, we considered three levels of cline variability: low (*σ_v_* = 0.2 and *σ_c_*= 0.5), moderate (*σ_v_* = 0.4 and *σ_c_* = 0.8), and high (*σ_v_* = 0.6 and *σ_c_* = 1.2) (for context, compare these to estimates of the same parameters across a series of empirical data sets in Firneno *et al*., 2023). We simulated 50 data sets for each level of cline variability. In each case, we sampled the cline parameters *v* and *c* from normal distributions (on the log_10_ and logit scale, respectively) with means of zero and standard deviations of *σ_v_*and *σ_c_*. Cline parameters were sampled for 100 loci per data set. We then sampled hybrid indexes for 50 hybrids per data set; these were drawn from a uniform distribution bounded by 0 and 1. We then computed the locus-specific ancestry for each locus *i* and individual *j* based on the cline parameters and hybrid index, *ϕ_ij_* = *H^vi^ /*(*H^vi^* + (1 *− H^vi^*)*e^ui^*), with *u_i_* = logit(*c_i_*)*v_i_*.

Local ancestry states for each locus and individual (*Z_ij_*) were then sampled from a binomial distribution with two draws using *ϕ_ij_* as the probability of ancestry from source population 1. In these initial simulations, we assumed fixed differences between source populations, such that ancestry was fully informative of state.

We then estimated cline parameters for each of the 150 data sets (50 replicates with each of three cline standard deviations). We analyzed the data using the standard hierarchical Bayesian genomic cline model in bgchm, and with two alternative models: (i) a non-hierarchical variant of the genomic cline model in bgchm with the prior cline standard deviation set to be relatively uninformative (*σ_v_* and *σ_v_* = 100) and the corresponding logit-logistic genomic cline model in HIest (version 2.0; Fitzpatrick, 2013a). The comparison with the non-hierarchical model was done to evaluate the effect of modelling the clines hierarchically versus not doing so. The comparison with HIest was chosen as this was the initial software developed to fit this form of genomic cine model (with a non-hierarchical model) and thus serves as a general check on the quality of our inference. Notably, only the hierarchical model provides a means to estimate the cline standard deviations and HIest requires fixed differences between parents (hence our focus on loci with fixed differences for this initial set of simulations). Genomic clines in HIest were fit using the L-BFGS-B algorithm. Models fit with bgchm used default HMC settings of 2000 iterations, including a 1000 iteration warmup, and no thinning. Four chains were run. For the hierarchical models, the priors on the standard deviations for *σ_v_* and *σ_c_*were normal with means of 0 and standard deviations of *σ*_0_ = 2. We used the known parental allele frequencies and hybrid indexes for all analyses (with the small caveat the parental allele frequencies of 0.001 and 0.999 were used rather than 0 and 1 to avoid problems with infinite probabilities during computation).

We conducted a second set of simulations to evaluate the effects of allele frequency differences between source populations and uncertainty in genotypes on the ability of bgchm to estimate genomic cline parameters. For this, we again simulated data using the logit-logistic genomic cline model as a generative model. Here, we considered only a case of intermediate variability in introgression across the genome, that is *σ_v_* 0.3 = and *σ_c_* = 0.7 (this is between the low and moderate variability cases considered for the first set of simulations). We simulated three levels of allele frequency differences between source populations: (i) fixed differences, (ii) SNPs with a minimum allele frequency difference of 0.5, and (iii) SNPs with a minimum allele frequency difference of 0.1. In each case, actual allele frequency differences for each SNP were sampled from a uniform distribution bounded by 1 and the specified lower bound (e.g., 0.5 or 0.1). Thus, allele frequency differences varied among loci (except in the case of all fixed differences), as would be expected for many empirical data sets. We simulated 50 data sets comprising 100 loci and 50 hybrids for each level of minimum allele frequency differences. Then, for each simulation, we generated an additional, complementary data set with uncertain genotypes. This was done as described above for the hybrid index and ancestry class coefficient analyses. Specifically, we again assumed a Poisson distributed number of reads per individual an locus (*λ* = 7) and 1% sequence error rate.

Next, we estimated genomic cline parameters for each of the 300 simulated data sets (50 replicates for each level of allele frequency differences and for genotypes known versus uncertain) using the hierarchical model from bgchm. We did not include the comparison with HIest as this program requires diagnostic allele frequency differences between source populations and we kept our focus on the hierarchical model to evaluate inferences of cline standard deviations. We used the default HMC settings of 2000 iterations, including a 1000 iteration warmup, and no thinning. Four chains were run. Priors on the standard deviations for *σ_v_* and *σ_c_* were normal with means of 0 and standard deviations of *σ*_0_ = 2. We again used the known parental allele frequencies and hybrid indexes for all analyses.

#### Genomic cline analyses of hybrid zones simulated with ***dfuse***

We then conducted a third set of simulations to examine the relationship between the genetic architecture of hybrid fitness and cline parameters, including both clines for individual loci and the cline parameter standard deviations. These simulations were not meant to be exhaustive but rather to complement existing simulation-based studies of genomic clines in the context of the genetics of isolation in hybrids and cline coupling (e.g., Gompert *et al*., 2012b; Firneno *et al*., 2023). Our purpose was to illustrate how different genetic architectures of hybrid fitness can leave different patterns in genomic clines and how these relate to patterns that might arise in the absence of selection.

Hybrid zones were simulated using dfuse (Lindtke & Buerkle, 2015). We described this software and model previously in the context of the simulations used to assess our hybrid index and ancestry class proportion models. In these individual-based simulations cline parameters are not strictly defined (i.e., it is not guaranteed that the patterns of introgression will conform precisely to the form specified by the genomic cline model nor are the parameters of such a model defined by the simulation conditions). Thus, we do not use these simulations to assess the accuracy of the bgcmh model *per se*, but rather to evaluate how cline parameter estimates are affected by the simulation conditions. Here, we assumed that hybrid fitness is determined by *N* underdominant loci, such that the fitness of an individual heterozygous for ancestry at *n* of the *N* loci is *w_j_* = (1 *− s*)*^n^*, where *s* is the selection coefficient (the underdominance model was added to dfuse in Firneno *et al*., 2023). We simulated ten replicate data sets under four different hybrid zone models. All simulations involved secondary contact, 15 demes for the hybrid zone, an adult carrying capacity of 100 individuals per deme, a migration rate of 0.05 between neighboring demes, and 5000 generations of evolution post secondary contact. We simulated hermaphroditic, diploid organisms each with one, 1 Morgan chromosome. We recorded ancestry at 251 evenly spaced loci along the chromosome of each individual. One set of simulations involved no selection (i.e., neutral evolution by drift and gene flow only). A second set assumed an oligogenic architecture of fitness with two underdominant loci with *s* = 0.3 at positions 25 cM and 75 cM along the chromosome (an individual heterozygous at both loci would have a relative fitness of 0.49). The third set of simulations considered a polygenic architecture with weak selection overall, specifically 50 underdominant loci distributed at even distances across the chromosome and with *s* = 0.005 per locus (an individual heterozygous at all 50 loci would have a relative fitness of 0.78). The last set of simulations was of strong polygenic selection, which again involved 50 evenly distributed underdominant loci but with *s* = 0.01 (an individual heterozygous at all 50 loci would have a relative fitness of 0.61).

We randomly sampled 100 individuals from each simulated hybrid zone for analysis. We assumed fixed differences between source populations at the 251 loci, such that ancestry was perfectly informative of genotype. Genomic cline parameters were estimated using the standard hierarchical model in bgchm. We used the known hybrid indexes and parental allele frequencies of 0.001 and 0.999. We again used the default HMC settings of four chains each comprising 2000 iterations including a 1000 iteration warmup, and no thinning. We set normal priors for *σ_v_* and *σ_c_* with means of 0 and standard deviations of *σ*_0_ = 2.

### Analysis of an example empirical data set

Lastly, to demonstrate possible usages of the genomic cline models in bgchm, we applied them to an empirical genetic data set from a hybrid zone in *Lycaeides* butterflies. The data set was originally published and analyzed in Chaturvedi *et al*.’s (2020). Two nominal species of *Lycaeides* butterflies, *L. idas* and *L. melissa*, occur throughout much of western North America with partially overlapping ranges (Nabokov, 1943; Gompert *et al*., 2010, 2014). These species differ on average in terms of the structure of the male genitalia (Nabokov, 1944; Gompert *et al*., 2010), aspects of wing pattern (Lucas *et al*., 2018), host plant species used, and voltinism (Gompert *et al*., 2013), but nonetheless have hybridized extensively (Gompert *et al*., 2010, 2012a; Nice *et al*., 2013; Gompert *et al*., 2014; Chaturvedi *et al*., 2020). An ancient, partially stabilized series of admixed populations occurs in the central Rocky mountains and Jackson Hole, which we refer to as Jackson Hole *Lycaeides*. These populations are the product of hybridization between *L. idas* and *L. melissa* about 14,000 years ago following the retreat of Pleistocene glaciers. More recently, Jackson Hole *Lycaeides* have come into secondary contact with *L. melissa* near the town of Dubois, WY (43.5623*^◦^*N, 109.6991*^◦^*W) where *L. melissa* feed on naturalized alfalfa (*Medicago sativa*) that grows along roadsides and that was introduced to North America about 250 years ago. This recent secondary contact has resulted in a contemporary hybrid zone (Chaturvedi *et al*., 2020; Zhang *et al*., 2023), which is the focus of our analyses here. Our goals here are to use bgchm to characterize the genomic composition of this hybrid zone in terms of hybrid indexes, ancestry class proportions and genomic cline parameters. We then specifically examine the extent to which clines differ on average between autosomes and the Z sex chromosome and as a function of features of the genome (gene and transposable element density).

We focus on a data set comprising the Dubois hybrid zone (*N* = 115) individuals, three populations representative of source Jackson Hole *Lycaeides* (set as source 0, *N* = 166), and two populations representative of source *L. melissa* populations (set as source 1, *N* = 117) (see Figure S1). We identified ancestry informative loci from a larger set of 39,193 SNPs generated from genotyping-by-sequencing data (see Chaturvedi *et al*., 2020 for details, including variant filtering and genotype inference). We specifically considered SNPs with an allele frequency difference of 0.3 or greater between our source populations; this yielded 500 ancestry informative SNPs (330 such SNPs on the 22 autosomes and 170 on the Z chromosome). We began by estimating hybrid indexes and ancestry class proportions based on this full data set with the est hi and est Q functions in bgchm. This was done using the known genotypes model with maximum likelihood estimates of parental allele frequencies derived from Bayesian point estimates of genotypes. We treated Z-linked SNPs in females as haploid. We used the default HMC conditions for these functions, that is four HMC chains with 2000 steps, including 1000 warmup iterations and no thinning. We summarized the posterior estimates of hybrid index and ancestry class proportions for each individual based on posterior medians and 90% CIs.

We next fit several genomic cline models to illustrate different ways cline estimates can be used to make inferences. First, as the primary analysis, we fit a single hierarchical genomic cline model using all 500 SNPs. Here, we used the hybrid indexes estimated from the full data set, maximum likelihood estimates of parental allele frequencies, and again treated Z-linked SNPs as haploid in females. We fit the model with the default HMC conditions–four chains each comprising 2000 iterations including a 1000 iteration warmup, and no thinning–with the prior mean for the cline standard deviations set to 0 and the prior standard deviations set to 2 (i.e., *σ_c_* and *σ_v_* were estimated from the data). We applied the sum-to-zero constraint to cline estimates from this analysis.

We then fit an additional pair of genomic cline models to directly ask whether patterns of introgression differed on average for autosomes versus the Z chromosome. For this, we used hybrid indexes estimated only from the autosomes. Genomic cline parameters for the autosomes and Z chromosome were then estimated separately, that is, in separate fits of genomic cline models. Here, not only did we estimate the cline standard deviations from the data (*σ_c_* and *σ_v_*), but also the mean (*µ_c_* and *µ_v_*). Because the hybrid indexes were based on the autosomal data, the expected means for the autosomal SNPs were *µ_c_* = 0 and *µ_v_* = 0. However, this was not true for the Z chromosome SNPs and the values of *µ_c_* and *µ_v_* for these SNPs thus indicate the extent and manner in which patterns of introgression deviate on average for Z-linked SNPs versus autosomes. With that said, the values of *µ_c_* and *µ_v_* for autosomes are not forced to be 0, and thus we based our inferences on the difference in *µ_c_* and *µ_v_* for Z for autosome SNPs (specifically, on the posterior distribution for such differences). These models were also fit the default HMC conditions, but with normal priors on *µ_c_* and *µ_v_*, both with means of 0 and standard deviations of *σ*_0_ = 2.

We conducted a final set of cline model fits to explicitly compare the variability of clines across autosomes versus the Z chromosome relative to the average introgression on autosomes versus the Z chromosome. For this we estimated hybrid indexes separately for autosomal and Z loci; we then fit hierarchical cline models for these sets of loci separately using the autosomal and Z-based hybrid indexes, respectively. We fixed cline means to 0 (as per the standard model) and estimated the cline standard deviations, *σ_c_* and *σ_v_*, which were the main focus of this analysis. This was again done with the standard HMC settings with the standard deviation of the normal prior on the cline standard deviations set to *σ*_0_ = 2. For all analyses we summarized the posterior estimates of cline parameters (*v* and *c*), hierarchical cline standard deviations and hierarchical cline means based on posterior medians and 90% CIs.

## Results

### Analyses of simulated data sets

#### Results for simulations of hybrid indexes and ancestry class proportions

Example graphical summaries of hybrid index (*H*) and interpopulation ancestry (*Q*_10_) estimates are shown in Figures 2A and 2B. In general performance was slightly better for hybrid index than interpopulation ancestry (Table S1 and Figure 2). Mean absolute error (MAE, the average deviation between true and estimated parameter values) increased with decreasing allele frequency differences, that is, with reduced information on ancestry in the genotypic data (Figure 2). However, even with allele frequency differences of 0.1, average MAEs were below 0.14 for *H* and 0.17 for *Q*_10_ (Table S1). Moreover, 90% credible intervals generally contained the true parameter value 90% of the time or more. Indeed, for the simulations with the greatest allele frequency differences (i.e., 1; fixed differences) the CIs appear to be conservative, with the true values of *H* and *Q*_10_ almost always falling within the 90% CIs. Inferences based on appropriately modelled uncertain genotypes were nearly as accurate as those based on known genotypes (Table S1 and Figure 2).

**Figure 1:**
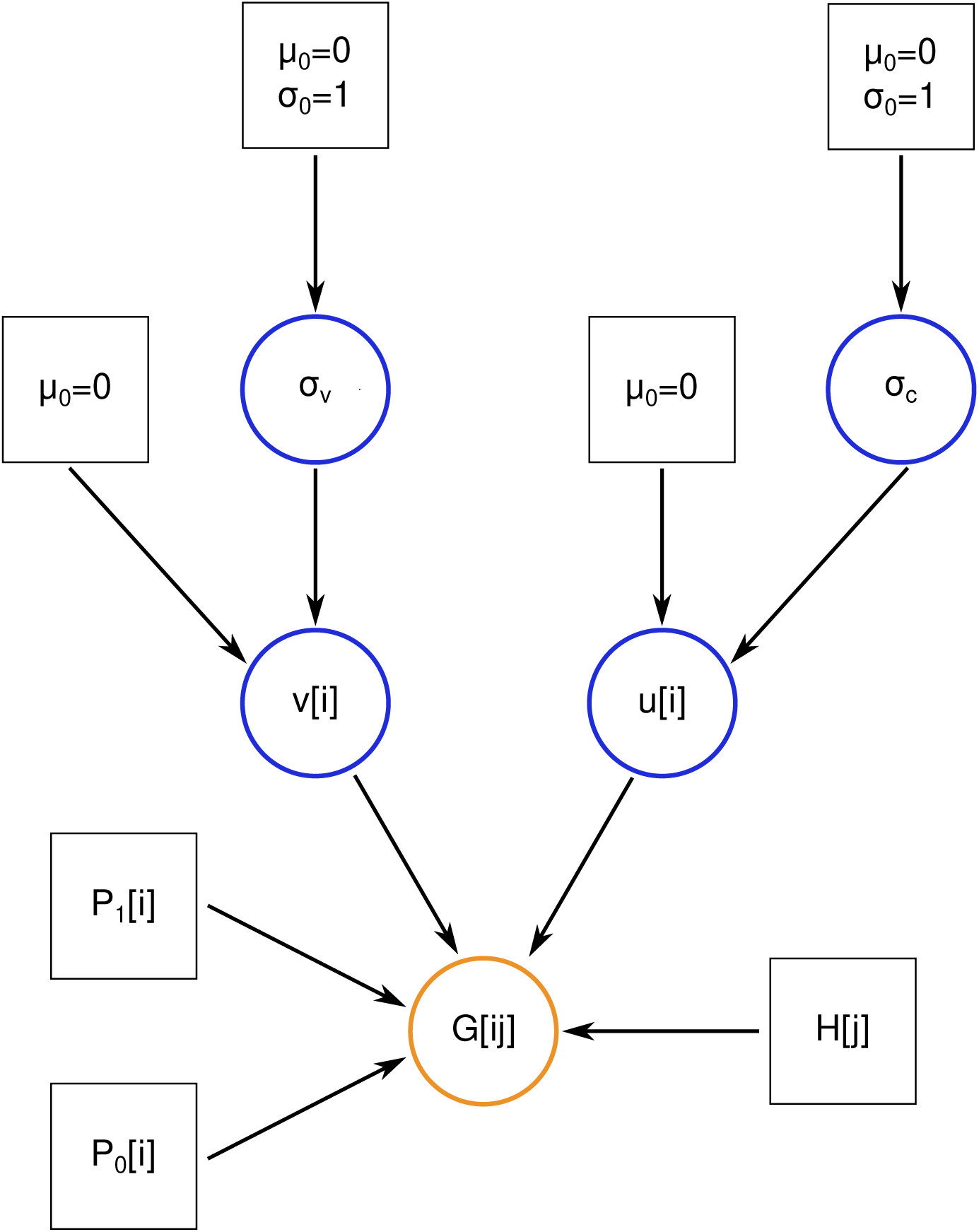
Directed graph summarizing the (standard) hierarchical Bayesian genomic cline model. Boxes and circles denote fixed and stochastic nodes, respectively, with the data node in orange and other stochastic nodes in blue. *G*[*i, j*] denotes the genetic data for locus *i* and individual *j*, that is, the known genotype or genotype likelihoods. *P*_0_[*i*] and *P*_1_[*i*] are the known (previously estimated) allele frequencies in parental source, reference populations. *H*[*j*] is the known (previously estimated) hybrid index for individual *j*. The stochastic nodes *v*[*i*] and *u*[*i*] are the cline parameters, with *v*[*i*] denoting the slope and *u*[*i*] = logit(center[*i*])*v*[*i*], where center[*i*] is the cline center. *σ_v_* and *σ_c_* denote the standard deviations of the normal priors on log(*v*[*i*]) and logit(center[*i*]). These describe variability in clines across the genome and are estimated from the data. The remaining fixed nodes denote means (*µ*_0_) and standard deviations (*σ*_0_) of higher-level normal priors.

**Figure 2:**
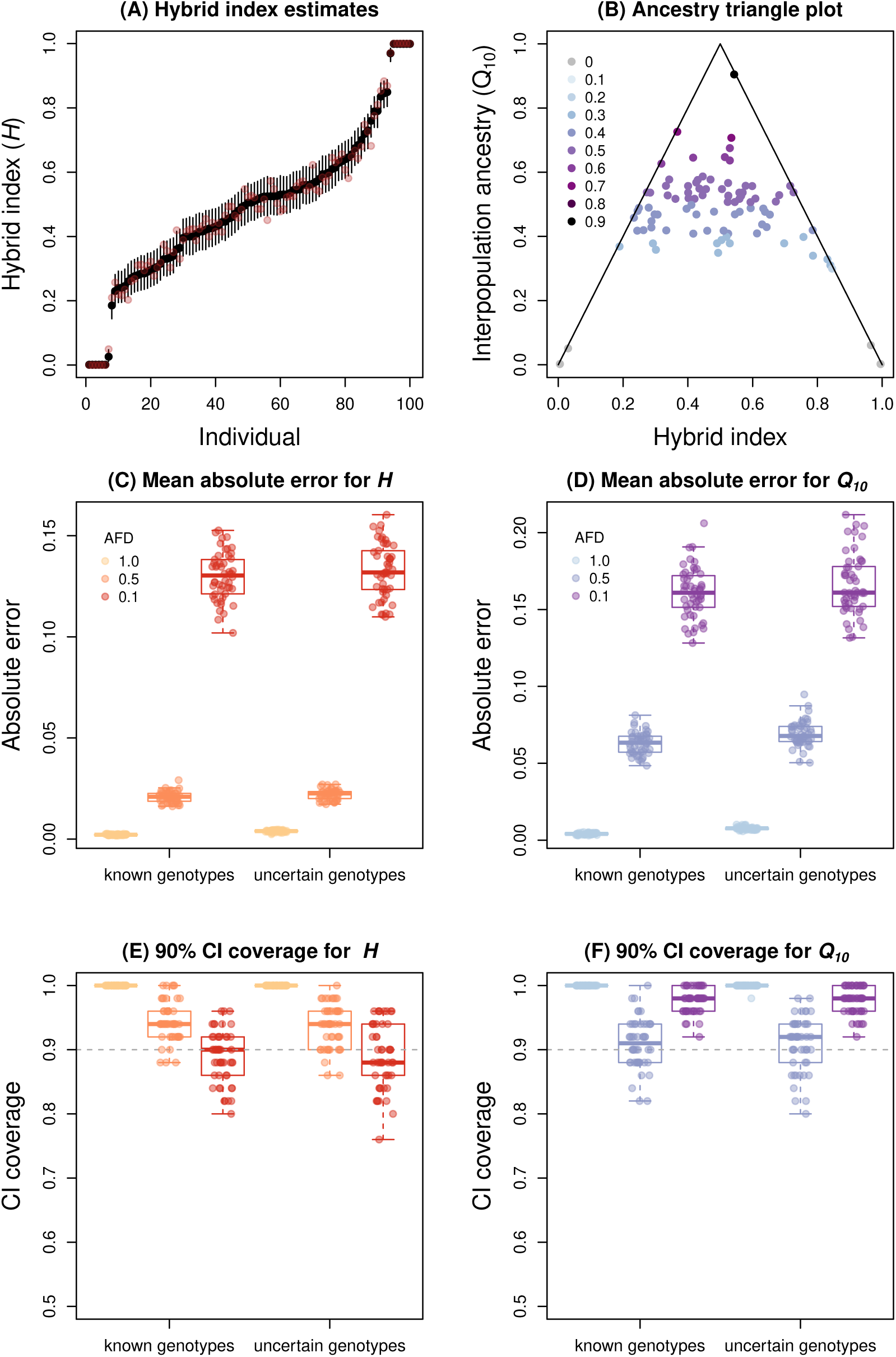
Example results and summary evaluation of model performance for estimating hybrid index (*H*) and interpopulation ancestry (*Q*_10_). Panel (A) shows point estimates of hybrid index (black points) and 90% credible intervals (CIs) (vertical lines) for 100 individuals. This is based on 100 loci with fixed differences between parental populations. Pink points show the true, simulated hybrid index values. The triangle plot in panel (B) shows interpopulation ancestry estimates (*Q*_10_) as a function of hybrid index (*H*) for the same simulated individuals. Point colors indicate true parameter values (in increments of 0.1) and lines (the triangle) denote maximum values of interpopualtion ancestry for a given hybrid index. Points on or near this line denote likely offspring with one non-hybrid parent. Panels (C)–(F) summarize model performance for 50 replicate simulations each with allele frequency differences (AFDs) between parents of 1.0, 0.5, or 0.1, and known or uncertain genotypes. Panels (C) and (D) summarize mean absolute error for estimates of *H* and *Q*_10_, respectively. Boxes indicate the median and 1st and 3rd quartiles of the distribution across replicate simulations, with whiskers extending up to 1.5*×* the interquartile range. The overlain points show metrics for individual replicates. Panels (E) and (F) similarly summarize the proportion of loci where the true parameter value is within the 90% CI of the Bayesian estimate for *H* (E) and *Q*_10_ (F). The horizontal dashed line denotes the expectation of 90% for a 90% CI.

#### Results from genomic cline analyses of simulated hybrid zones

Example clines for loci with fixed differences and with low (*σ_v_*= 0.2 and *σ_c_* = 0.5), moderate (*σ_v_* = 0.4 and *σ_c_* = 0.8), and high (*σ_v_* = 0.6 and *σ_c_*= 1.2) variability in introgression across the genome are shown in Figure 3A. Under these conditions, estimated cline standard deviations were highly correlated with the true cline standard deviations, with Pearson correlations of 0.97 (95% confidence interval = 0.96-0.98) and 0.97 (95% confidence interval = 0.95-0.98) for *σ_v_* and *σ_c_*, respectively (Figure 3B,C). With that said, when cline variability variability was high, variation was somewhat underestimated, such that the mean estimates of *σ_v_* and *σ_c_* for the highest variability case were 0.50 and 1.09 compared to the true values of 0.6 and 1.2. Such a bias was not apparent for the low variability simulations (mean *σ_v_* = 0.20 and mean *σ_c_* = 0.47, compared to true values of 0.2 and 0.5).

**Figure 3:**
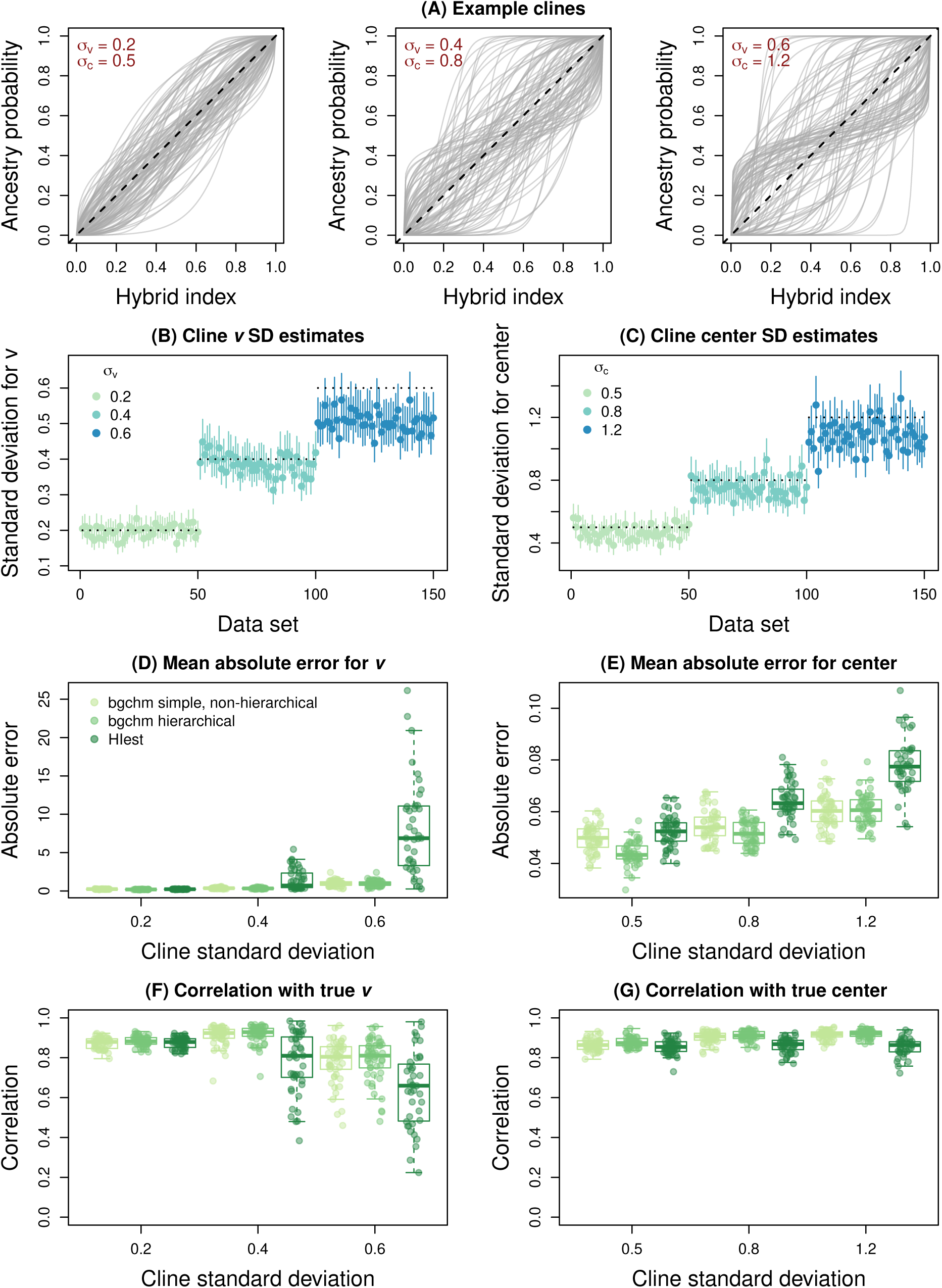
Summary of genomic cline variability and the effect of such variability on cline inference. Panel (A) shows simulated genomic clines with low (*σ_v_* = 0.2 and *σ_c_* = 0.5), moderate (*σ_v_* = 0.4 and *σ_c_* = 0.8), and high (*σ_v_*= 0.6 and *σ_c_* = 1.2) variability in introgression. Each gray line is the cline for a locus and gives the probability of ancestry from source 1 as a function of hybrid index (the overall proportion of the genome from source 1). The null expectation if introgression does not vary across the genome is given by the dashed black line. Estimates of cline standard deviations for slope, *σ_v_*, and center, *σ_c_*, are shown in panels (B) and (C), respectively. Here, point estimates and 90% credible intervals (CIs) are depicted with points and vertical lines. Horizontal dotted lines give the true value used for each simulation. Performance, in terms of estimating cline slopes (*v*) and centers, is summarized based on mean absolute error in panels (D) and (E) and in terms of the correlation between true and estimated parameter values in panels (F) and (G). Errors and Pearson correlations were computed based on parameter point estimates (posterior medians) and are summarized across replicate simulations with boxplots. Boxes indicate the median and 1^st^ and 3rd quartiles of the distribution across replicate simulations, with whiskers extending up to 1.5*×* the interquartile range. The overlain points show metrics for individual replicates. Performance of bgchm using a simple non-hierarchical model and a hierarchical model are shown, as are results from HIest (for HIest, cases where the algorithm failed are excluded).

With regard to individual cline parameters, mean absolute error was generally higher when clines were more variable, and likewise, correlations between true and estimated values declined (more so for slope than center) (Table S2, Figure 3). In general, bgchm outperformed HIest, especially when cline variability was high. The hierarchical and non-hierarchical models performed similarly, but with slightly better performance in terms of error and correlations with true parameter values for the hierarchical model, especially when cline variance was low (Table S2). As expected, our results also suggest that, relative to the non-hierarchical model with weakly informative priors, the hierarchical model is conservative in the sense that it induces some shrinkage towards zero into the parameter estimates (see Table S3). Furthermore, *σ_v_* and *σ_c_* can only be estimated as model parameters in the hierarchical model.

We next considered the effects of source allele frequency differences and genotype uncertainty on estimates of genomic cline parameters with the hierarchical model in bgchm. We found that cline standard deviation estimates (*σ_v_* and *σ_c_*) were most accurate for fixed differences, and became progressively less accurate with low levels of allele frequency differences (Figure 4A,B). When the minimum allele frequency difference between source populations was 0.1, there was a tendency to overestimate the variability in cline slopes (true *σ_v_* = 0.3, mean point estimate for known genotypes = 0.39) and underestimate the variability in cline centers (true *σ_c_* = 0.7, mean point estimate for known genotypes = 0.58). Uncertainty in genotypes had little effect on estimates of cline standard deviations (Figure 4A,B). Similarly, cline parameter estimates were most accurate in terms of both mean absolute error and the correlation with true parameter values when allele frequency differences were high, and were less accurate when they were low (e.g., 0.1; Table S4 and Figure 4). Uncertainty in genotypes tended to further decrease the accuracy of estimates, but only to a minor extent (see Table S4 and Figure 4). Moreover, the average proportion of loci where the true parameter value was contained in the 90% CIs was only weakly affected by allele frequency differences or genotype uncertainty suggesting that the uncertainty in clines caused by weak genetic differentiation between sources is mostly captured by the uncertainty in parameter estimates (Table S4). With that said, there was slight tendency overall to underestimate cline uncertainty (i.e., between 80 and 88 percent of the 90% CIs contained the true value relative to the expectation of 90 percent).

**Figure 4:**
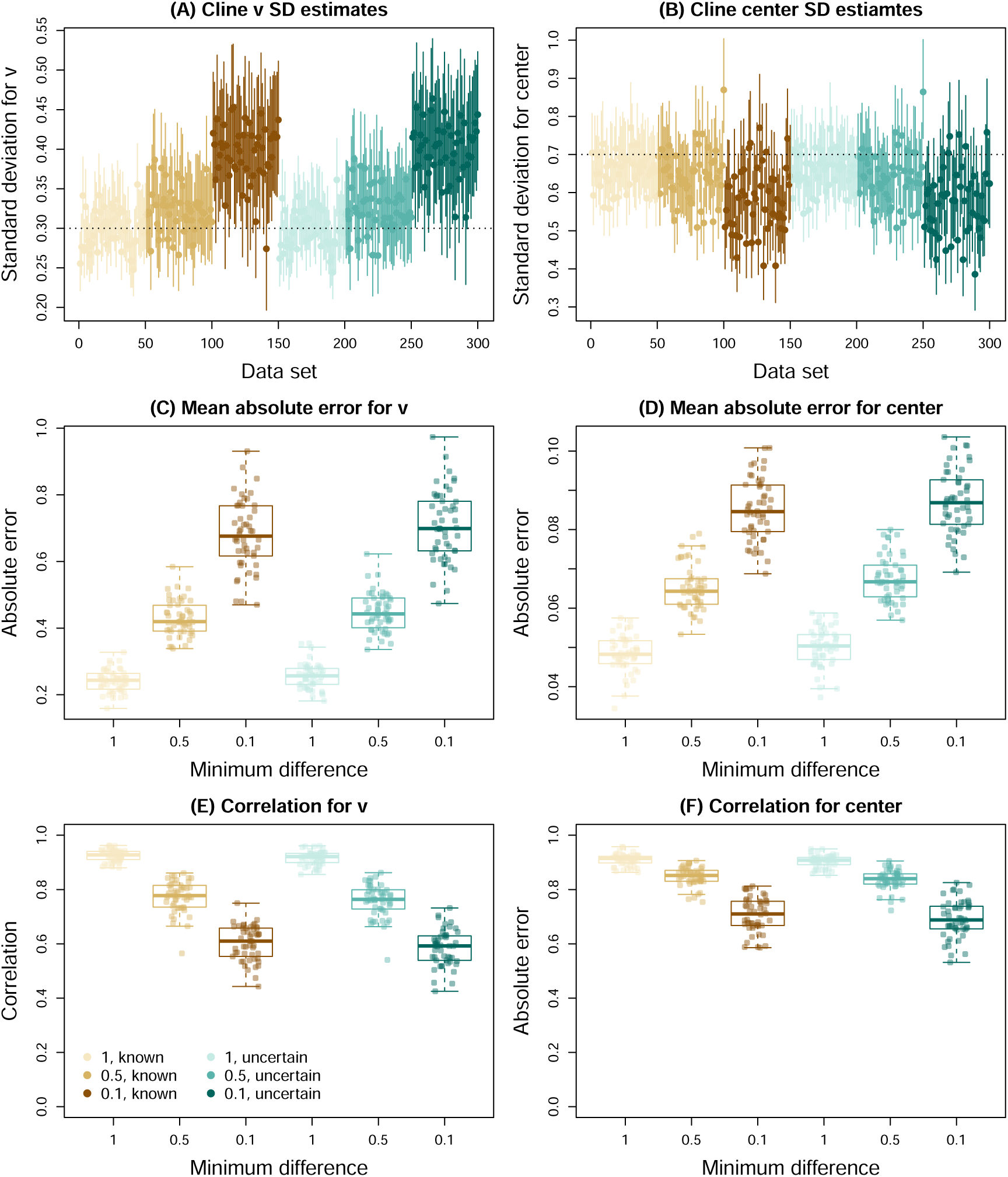
Summary of the effects of source allele frequency differences and genotype uncertainty on genomic cline inference. All panels show results based on minimum source allele frequency differences of 1, 0.5 and 0.1 and with or without uncertainty in genotypes as indicated by the colors and associated legend. Panels (A) and (B) provide estimates of cline standard deviations for slope, *σ_v_*, and center, *σ_c_*, respectively. Points and vertical lines depict point estimates and 90% credible intervals (CIs) Horizontal dotted lines give the true value used for *σ_v_* (A) and *σ_c_* (B). Model performance for genomic cline parameters (*v* and *c*) is summarized based on mean absolute error in panels (C) and (D) and based on the correlation between true and estimated parameter values in panels (E) and (F). Errors and Pearson correlations were computed based on parameter point estimates (posterior medians) and are summarized across replicate simulations with boxplots. Boxes indicate the median and 1^st^ and 3rd quartiles of the distribution across replicate simulations, with whiskers extending up to 1.5*×* the interquartile range. The overlain points show metrics for individual replicates.

#### Results from genomic cline analyses of hybrid zones simulated with ***dfuse***

Our final analysis of simulated hybrid zones involved various genetic architectures for hybrid fitness with dfuse. Overall, stronger selection (oligenic or strong polygenic) resulted in a steeper geographic clines in hybrid indexes across the hybrid zones (Figure 5). However, all four sets of conditions resulted in fairly similar numbers of loci with credible deviations from null expectations for cline slopes (*v*) and centers (*c*) (Table S5 and Figure 5). We observed notable variation in cline standard deviations across simulated data sets, with a trend towards larger slope variances (*σ_v_*) for oligogenic selection (Figure 5I).

**Figure 5:**
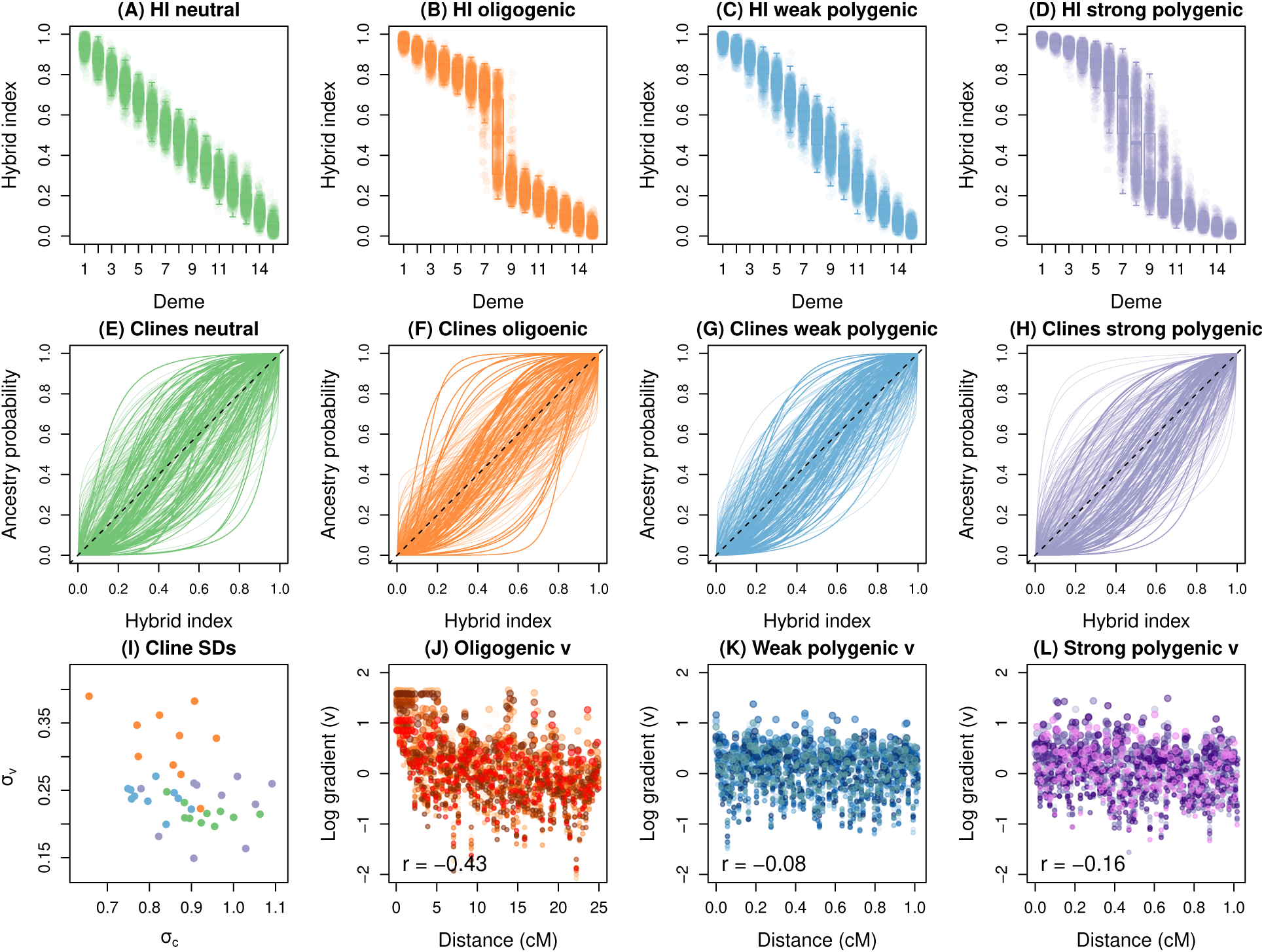
Summary of genomic cline analysis of hybrid zone simulations with alternative genetic architectures for hybrid fitness. Results are shown for neutral secondary contact, oligenic selection, weak polygenic selection and strong polygenic selections (see main text for details). Panels (A–D) show the distribution of hybrid indexes (HI) across demes and across 10 replicate simulations for each set of conditions. Boxes indicate the median and 1st and 3rd quartiles of the hybrid index distribution for each deme, with whiskers extending up to 1.5*×* the interquartile range. The overlain points denote individual hybrid indexes. Panels (E–H) show genomic clines from 100 representative loci for each set of conditions. Each colored line is the cline for a locus and gives the probability of ancestry from source 1 as a function of hybrid index. The null expectation if introgression does not vary across the genome is given by the dashed black line. Point estimates of cline standard deviations (SDs) are shown in panel (I). Here, conditions are colored in accordance with panels (A–H). Panels (J), (K) and (L) show the relationships between the distance (in cM) a marker locus is from a selected locus and the log of the cline gradient or slope (*v*). This is only shown for the three sets of conditions with selection. Points are colored to indicate distinct replicate simulations and the Pearson correlation between distance and log(*v*) is reported.

Despite similar numbers of loci with clines deviating from null expectations, we did find patterns of cline variation consistent with the effects of selection. Specifically, for oligogenic selection and strong polygenic selection there was a significant (all *P <* 0.05) negative correlation between the log of *v* and the distance a marker locus was from an underdominant locus (Pearson correlations ranged from -0.34 to -0.49 for oligogenic selection and -0.11 to -0.23 for strong polygenic selection; Table S6 and Figure 5). Negative correlations were also observed for weak polygenic selection (range = -0.04 to -0.11), but these were not significantly different from 0 (all *P >* 0.05) (Table S6 and Figure 5). No underdominant loci were present in the neutral simulations, thus, as expected, we found small and non-significant (and mostly positive) correlations between cline slopes and the locations used for underdominant loci in the polygenic simulations (range = -0.02 to 0.06) (Table S6); this demonstrates that large negative correlations do not arise inherently in the absence of selection.

Interestingly, we detected positive correlations between the absolute value of logit cline centers and the location of underdominant loci in the oligogenic simulations and most of the strong polygenic simulations (positive in all ten of the latter, but significantly greater than 0 with *P <* 0.05 for eight of the simulations; Table S7). A similar but non-significant pattern was documented for weak polygenic selection and no such pattern was found for neutral simulations (again based on the locations of underdominant loci in polygenic simulations). Thus, at least with stronger selection, simulated SNPs near underdominant loci have steeper cline slopes (larger, positive values of *v*) but also cline centers closer to the genome-wide null expectation, suggesting that selection resulted in steeper clines but more constrained (coincident) centers.

### Analysis of an example empirical data set

We estimated hybrid indexes, ancestry class proportions, and genomic cline parameters for 500 ancestry-informative SNPs in a *Lycaedies* butterfly hybrid zone (Figure 6). Estimates of hybrid indexes were generally precise (mean width of the 90% CIs = 0.061) and spanned the full range from only ancestry from source population 0 (i.e., Jackson Hole *Lycaeides*) to only ancestry from source population 1 (i.e., *L. melissa*) (Figure 6B). Ancestry class proportion estimates suggest a wide range of genome compositions in hybrids, including some individuals with near maximal interpopulation ancestry for their hybrid indexes (i.e., individuals with one or more non-admixed parents, that is F1s or backcrosses) and individuals where both parents were likely themselves hybrids (i.e., individuals with lower levels of interpopulation ancestry given their hybrid indexes; Figure 6C).

**Figure 6:**
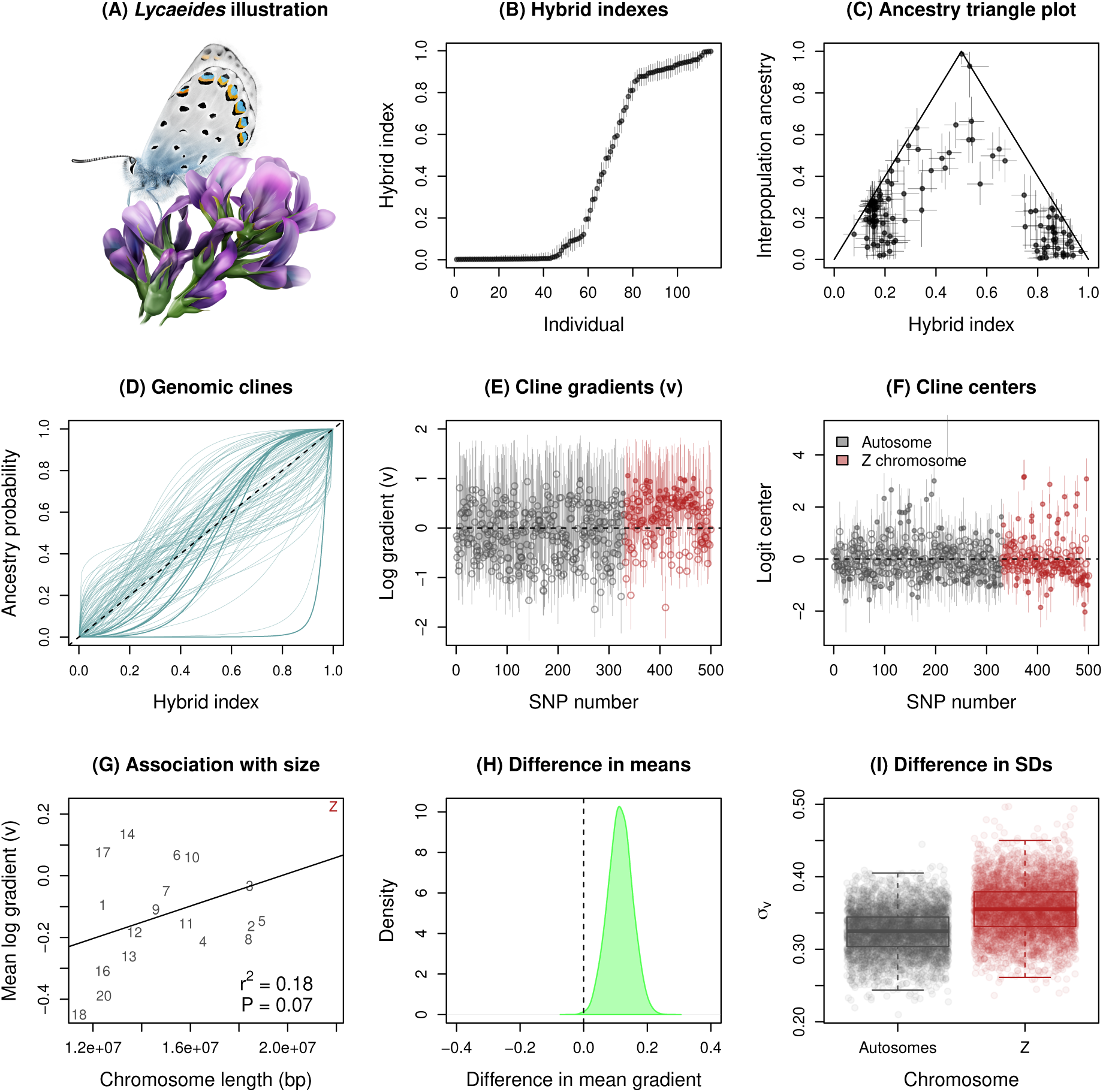
Summary of key results from an example analysis with *Lycaeides* butterflies. An illustration of a *Lycaeides* butterfly from the Dubois hybrid zone is shown in panel (A). Panel (B) gives point estimates (points) and 90% credible intervals (CIs) (vertical lines) for hybrid index based on the combined autosomal and Z chromosome data. Panel (C) shows interpopulation ancestry estimates (*Q*_10_) as a function of hybrid index (*H*) for the same hybrid zone butterflies. Point estimates and 90% CIs (vertical and horizontal lines) are given. Genomic clines for 100 representative loci are shown in panel (D). Each line denotes the probability of *L. melissa* ancestry for a locus as a function of hybrid index (the overall proportion of an individual’s genome with *L. melissa* ancestry). Darker and thicker lines are used for loci with credible deviations from genome-average ancestry (90% CIs for cline gradient of center not overlapping null expectations). The diagonal, dashed line shows the null 1:1 expectation for locus-specific ancestry probabilities as a function of hybrid index. Panels (E) and (F) display estiamtes of the log cline gradient (log of *v*) and logit cline center for each of the 500 ancestry-informative SNPs. Point estimates and 90% CIs (vertical lines) are displayed, with open points used for cases where the 90% CIs do not exclude values less than 0 (E) or do not exclude 0 (F). The null expectation value of 0 (on the log or logit scale) is shown for each panel with a horizontal dashed line. Panel (G) shows the relationship between chromosome size (length in base pairs, bps) the the mean log gradient for the 20 chromosomes with more than five ancestry informative SNPs. Chromosome numbers (or Z) are given, along with the best fit line from a linear regression; the model *r*^2^ and *P* -value are reported. Panel (H) gives the difference in mean log gradient between the Z chromosome and autosomes for cline models where Z and autosomal SNPs were analyzed separately and where the means were not set to zero but estimated from the data. Both models used autosomal estimates of hybrid indexes. The posterior density for the difference is shown, along with a vertical line for the null expectations. The posterior probability that the mean for Z loci exceeds the mean for autosomes was *>* 0.99. Panel (I) shows the posterior distributions for the standard deviation in log cline gradients for autosomes and the Z chromosome. Here, autosomal and Z SNPs were analyzed separately and with hybrid indexes inferred from autosomal and Z SNPs, respectively. Boxes indicate the median and 1st and 3rd quartiles of the posterior distribution, with whiskers extending up to 1.5*×* the interquartile range. The overlain points show 4000 parameter value samples from the posterior. The posterior probability that the variance for the Z SNPs exceeds the variance for the autosomal SNPs was 0.75.

Genomic cline analysis of all 500 SNPs detected substantial genome-wide variation in introgression (Figure 6D–F). Overall, patterns of introgression deviated from null expectations based on genome-average admixture for 218 out of the 500 loci (Table S8). This includes 40 loci with credibly steeper clines than null expectations (*v >* 1), of which 39 were on the Z chromosome. This is a significant enrichment of steep clines on the Z chromosome (randomization test, 1000 randomizations, expected = 13.7, *P* = 0.001). We detected 48 loci with credible excesses in Jackson Hole *Lycaeides* (*c >* 0.5) or *L. melissa* (*c <* 0.5) ancestry, with enrichments of both types of excesses on the Z chromosome (randomization tests, 1000 randomizations each; *c >* 0.5, Z observed = 23, Z expected = 16.4, *P* = 0.022; *c <* 0.5, Z observed = 37, Z expected = 23.8, *P* = 0.001).

In general, variability in introgression among loci can reflect the joint effects of selection and genetic drift. A role for selection predicts associations between cline parameters and genomic features, such as chromosome size and genomic content (e.g., Schumer *et al*., 2018; Chaturvedi *et al*., 2020). Along these lines, we found a modest and marginally significant positive association between chromosome size and mean log cline slope or gradient (*v*) when considering the subset of chromosomes with at least five ancestry informative SNPs (linear regression, df = 17, *β* = 2.6 *×*10*^−^*8, s.e. = 1.4 *×*10*^−^*8, *r*^2^ = 0.18, model *P* = 0.017). This would be expected if loci on larger chromosomes were affected on average more by indirect selection because of a lower rate of recombination per base pair (and thus higher average linkage disequilibrium). We found no evidence of steeper clines in or near (within 1 kilobase) genes (randomization test, 1000 randomizations, *P* = 0.792), but did find evidence of significantly steeper clines in or near annotated transposable elements (randomization test, 1000 randomizations, *P* = 0.011). Together these results suggest some role for selection in clinal patterns, and highlight different patterns of introgression for autosomes and the Z chromosome. We followed up on this latter possibility with formal analyses comparing these sets of chromosomes.

In cline models based on atuosomal hybrid indexes, we found credibly steeper clines on average for the Z chromosome than for autosomes (posterior probability *µ_v_* for Z was greater than *µ_v_* = 0.999 for autosomes, estimate of difference = 0.118, 90% CIs =0.055-0.181; Figure 6H). This is consistent with stronger selection in hybrids on the Z chromosome, especially as drift has a much more pronounced effect on cline centers than slopes in the absence of spatial structure (Gompert *et al*., 2012b). When considering cline slopes inferred from fully independent analyses of autosomes and Z loci (for hybrid indexes and clines), we found a trend towards more variability of introgression on the Z relative to average introgression on the Z (*σ_v_* = 0.355, 90% CI = 0.296-0.415) versus variability of introgression on the autosomes relative to average introgression on autosomes (*σ_v_* = 0.325, 90% CI = 0.273-0.374), but there was sufficient uncertainty in both parameters to preclude strong confidence in the difference suggested by this trend (posterior probability Z *>* autosomes = 0.748, see Figure 6I). Still, taken together these results point to a special role for the Z sex chromosome in speciation in *Lycaeides* butterflies (consistent with Chaturvedi *et al*., 2020)

## Discussion

Genomic analyses of hybrid zones provide unique and powerful insights into the nature and basis of species boundaries and the ecological and evolutionary consequences of hybridization (Harrison & Larson, 2014; Gompert *et al*., 2017). Here, we described, demonstrated and assessed bgchm, a new R package designed to facilitate genomic analyses of hybrid zones. This R package combines methods and models for Bayesian inference hybrid indexes, ancestry class proportions, and genomic clines (and also geographic clines, see the OSM, Table S9 and Figure S3) with HMC. We showed that bgchm provides accurate estimates of the relevant model parameters under a variety of conditions, and especially when the genetic loci are highly informative of ancestry (i.e., when the allele frequency differences between source populations are not too small). This even includes reasonably robust estimates of the variability of clines across the genome via inference of cline standard deviations, which have not been the focus of previous models and methods. The models presented also allow for inference with uncertainty in genotypes, and we showed that at least with modest sequencing coverage this has minimal effect on the accuracy of inferences. Finally, we found that under most conditions true uncertainty in parameters was accurately estimated, although in some cases credible intervals were overly conservative (e.g., hybrid indexes with fixed differences between parents) or too narrow (e.g., genomic cline parameters in some cases).

Our results from simulated and empirical data sets build on our existing understanding of how evolutionary processes interact to affect patterns of introgression in hybrid zones (e.g., Endler, 1977; Barton & Hewitt, 1985; Gompert *et al*., 2012b; Harrison & Larson, 2016; Gompert *et al*., 2017; McFarlane *et al*., 2021). For example, when hybrid fitness has a simple genetic architecture, loci residing in genomic regions proximate to causal variants affecting hybrid fitness had exceptional genomic cline parameters, consistent with Gompert *et al*. (2012b). The effects of selection on individual genomic cline parameters were less pronounced for weaker and more polygenic selection, though some signals remained in terms of cline parameters varying as a function of distance from causal variants in simulations and differences among classes of loci (those near transposable elements or on the Z sex chromosome versus autosomes) for the *Lycaeides* hybrid zone. This suggests that when the genetic architecture of hybrid fitness is polygenic, it is probably more informative to focus on such higher level contrasts, including cline standard deviations (which can sometimes be related to cline coupling, see, e.g., Firneno *et al*., 2023), rather than so-called individual outlier loci. It is also critical to recall that selection is not required for introgression to vary across the genome and for loci to deviate from null patterns of introgression based on genome-wide admixture, as illustrated by our simulations of neutral secondary contact. Indeed, selection can either increase or decrease the variation in introgression across the genome, with the former expected for simple genetic architectures and the latter expected for coupled clines when many loci contribute to reproductive isolation (Barton, 1983; Firneno *et al*., 2023). Thus, additional information beyond deviations from genome-average introgression is required to infer processes from patterns in hybrid zones; we expand on this topic in the section below on “Conclusions and future directions”.

### Comparison with other software

Several computer programs exist for genetic analyses of hybrid zones, and thus it is worth considering how this newly introduced R package, bgchm, fits in with existing software. To our knowledge, four main programs are currently available for estimating genomic clines. The earliest of these was introgress (Gompert & Buerkle, 2010), which adopts a multinomial likelihood-based approach to estimate genomic clines for multilocus genotypic data (Gompert & Buerkle, 2009, 2010). This program does not consider ancestry but is unique in separately modelling introgression of homozygous versus heterozygous genotypes. The original bgc (Gompert & Buerkle, 2012) fits Bayesian genomic clines in ancestry using a hierarchical model and a polynomial function for clines adapted from Szymura & Barton’s (1986). This software has many similarities with our new bgc, including the basic hierarchical modelling approach and the ability to work with genotype uncertainty. However, bgc is less modular (all loci must be fit together) and uses traditional Markov chain Monte Carlo, which exhibits notably poorer mixing. These features make bgc less well-suited for genome-scale data and for estimating cline standard deviations (these tend to mix especially poorly and are generally treated as nuisance parameters). HIest features multiple cline models and approaches to model fitting, but takes a non-hierarchical approach and assumes fixed differences between source populations (Fitzpatrick, 2012). Finally, the recently released gghybrid (Bailey, 2022), shares many aspects with bgchm, including the Bayesian approach and use of the logit-logistic cline model. The key differences are that gghybrid fits a non-hierarchical model and only considers the case of modelling known genotypes. Furthermore, gghybrid does not use HMC for inference. In terms of speed, the original bgc is by far the slowest program, especially with large data sets, whereas the likelihood based approaches tend to be the fastest. We have not conducted a detailed comparison of gghybrid and bgchm, and this is slightly complicated by the fact that fewer MCMC steps are required to obtain a high effective sample size with HMC, but both programs make it practical to analyze very large data sets. For bgchm, the total runtime largely depends on the extent to which cline fitting is done in parallel after the cline standard deviation parameters have been estimated. With robust computational resources (i.e., a single compute node with *∼*48 CPUs and multi-threading), we have been able to successfully fit clines for millions of SNPs in a few days of human time.

Further, existing programs differ in terms of the set of features included. The original bgc was a standalone program that only included the genomic cline model but did estimate hybrid indexes as part of this model. In contrast, introgress, HIest, and gghybrid include additional functions for estimating hybrid indexes and (for the former two) for estimating genotype-based metrics similar to ancestry class proportions. bgchm also includes models of hybrid index inference and includes a unique model for true ancestry class proportions (this is similar to the Q model in entropy, but with source populations designated *a priori* ; Gompert *et al*., 2014; Shastry *et al*., 2021). Finally, while several computer programs, including Cfit (Gay *et al*., 2008) and hzar (Derryberry *et al*., 2014), which uses a Bayesian approach, exist for inference of geographic cline parameters, bgchm is unique in including the option to fit hierarchical models for geographic and genomic clines in a single program (the geographic cline models are described in the OSM).

Lastly, likelihood based approaches for estimating hybrid index and interpopulation ancestry exist in introgress Gompert & Buerkle, 2010 and HIest (Fitzpatrick, 2012), and hybrid indexes can be inferred in several programs using either likelihood or Bayesian methods (e.g., introgress, HIest, bgc and gghybrid; Gompert & Buerkle, 2012; Bailey, 2022). Moreover, interpopulation ancestry and admixture proportions, which are analogous to hybrid indexes with two source populations, can be jointly inferred in entropy (Gompert *et al*., 2014; Shastry *et al*., 2021). Finally, similar to the other R-based software packages, bgchm includes various functions for plotting results.

### Conclusions and future directions

Our use of HMC, and specifically the NUTS algorithm from Stan, results in more rapid and robust Bayesian inference of genomic clines than was possible with the original bgc program. However, analyses of very large data sets, or of many replicate hybrid zones, can still require substantial time or computational resources. One possible way to overcome this limitation is to replace the current HMC approach with an approximation of the posterior through variational inference (Kucukelbir *et al*., 2017). Variational inference is supported by Stan and allows for automatic approximation of the posterior distribution. This can increase the speed of model fitting by orders of magnitude (Kucukelbir *et al*., 2017). However, it can also come at a cost in terms of accuracy, and the reliability of variational inference for genomic cline models remains to be evaluated. We see additional potential for increases in speed, and potentially accuracy, by fitting cline models for ancestry blocks (as identified in models for local ancestry inference, e.g., Sankararaman *et al*., 2008; Browning *et al*., 2023) rather than for genotypes or ancestry at individual loci. This could reduce the number of independent genomic regions or loci required for analysis and simultaneously overcome limitations that arise from low ancestry information for subsets of loci. We intend to evaluate both variational approximations and ancestry-block based analyses in a future publication.

Finally, hybrid indexes, ancestry class proportions and genomic clines provide summaries of patterns of introgression, but connecting such genomic patterns to ecological and evolutionary processes remains difficult (McFarlane *et al*., 2021). With certain assumptions or information, especially about dispersal, geographic patterns of introgression can be directly related to process-based parameters, such as the average intensity of selection against hybrids (e.g., Barton & Hewitt, 1985; Szymura & Barton, 1986; Mallet *et al*., 1990). However, this is less true for genomic clines, as these are always relative to overall admixture and thus not absolute metrics of introgression (this is also an advantage as they are less dependent on the geography of a hybrid zone). We think a valuable area for future research is to test whether the combined information from hybrid indexes, ancestry class proportions, genomic and geographic clines, as well as patterns of linkage disequilibrium in hybrid zones, could be used to reliably infer demographic and evolutionary processes governing hybrid zones, at least for a subset of clear, alternative models. This could be done using approximate Bayesian computation or with neural networks, both of which are suitable for combining information across heterogeneous data types (Sisson *et al*., 2018; Gehara *et al*., 2020; Yang *et al*., 2022). Convolutional neural networks, which have recently shown great general promise in population genomics (Flagel *et al*., 2019; Torada *et al*., 2019; Smith *et al*., 2023), could be particularly useful for mapping such disparate data information sources to generative processes that emit identifiable signals. We think that this gap between pattern and process is an important area for future work to address and we hope to contribute to doing so in future work.

### Author contributions

Conceived the study: ZG and CAB, initial software development: ZG, data analysis: ZG, initial draft of the manuscript: ZG, revised and edited the software code, help files and manuscript: ZG, DAD and CAB.

## Acknowledgments

We thank Rozenn Pineau, Alia Donley, Anthony Reis and Bhagya Amarasinghe for their help with testing the bgchm R package. This work was supported by NSF grant DEB 1844941 to ZG. Support and resources from the Center for High Performance Computing at the University of Utah are gratefully acknowledged.

## Data availability

Simulated data sets are available from Dryad (DOI pending). DNA sequence data for the butterfly hybrid zone are available from the NCBI SRA (PRJNA577236 and PRJNA432816).

## Code availability

The source code for bgchm can be downloaded and installed from GitHub (https://github. com/zgompert/bgc-hm). Additional scripts used for simulations and analyses in this article are available from a second GitHub repository (https://github.com/zgompert/bgchm_ test).

## Supplemental Methods and Results

### Geographic cline model

A variety of hybrid zone models predict sigmoidal allele frequency clines such that *p* = 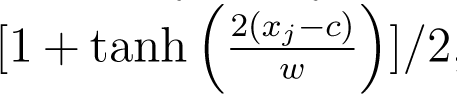, where *w* is the cline width, *c* is the cline center, and *x* is the location of deme *j* (e.g., Szymura & Barton, 1991; Barton *et al*., 1993). This includes cases where clines are neutral (e.g., result from recent secondary contact and are not maintained by selection) and cases where clines are maintained by the oppossing processes of dispersal and various forms of endogeneous or exogenous selection (Barton & Hewitt, 1985; Barton *et al*., 1993). When many loci contribute to hybrid fitness, clines can be coupled such that each experiences, indirectly (via linkage disequilibrium), some of the selection caused by the effects of the other loci (this applies even to neutral loci) (Barton, 1983). When this occurs, clines are expected to retain the sigmoidal form in the center of the hybrid zone (with a width dictated by direct and indirect selection), but exhibit a distinct pattern of exponential decay in allele frequency away from the hybrid zone center (Barton, 1983). Thus, the sigmoidal function applies to both single locus and multi-locus clines, but only away from the edges of the hybrid zone. This central, sigmoidal, portion of the hybrid zone can be described by a linear function on a logit scale, that is logit(*p_ij_*) = *c_i_* +*β_i_x_j_* (Szymura & Barton, 1991; Barton *et al*., 1993). Here, *c_i_* and *β_i_*are the center (intercept on the logit scale) and slope of the cline for locus *i* and *x_j_* is the (centered) geographic coordinate for deme *j*. The cline width on the natural scale (*w_i_*) is related to the slope as 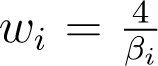. We work with this logit-linear geographic cline function.

We specifically assume the logit-linear function above describes the allele frequency cline over some geographic region, specifically the geographic region where allele frequencies vary between a defined minimum and maximum value (i.e., away from the edges of the cline or hybrid zone). In bgchm, this value can be set by the user, but we use -2 to 2 on the logit scale by default, *∼*0.12 to *∼*0.88 on the natural scale (as in Kruuk *et al*., 1999). We thus assume that the likelihood of logit(*p_ij_*) (logit allele frequency for locus *i* and population *j*) is logit(*p_ij_*) *∼* normal(*c_i_*+ *β_i_x_j_, σ_ɛ_*). In this model *σ_ɛ_* is a standard deviation parameter describing the residual error.

Our main interest is in estimating the slopes (and associated cline widths). We do this using a hierarchical model, which also allows us to estimate the variability in slopes (and thus cline widths) across the genome (see the main text for a discussion of the general benefits of hierarchical modelling). We thus place a normal prior on the slopes, *β_i_ ∼* normal(*µ_β_, σ_β_*), where *µ_β_* and *σ_β_* describe the mean and standard deviation of slopes and are estimated from the data (this is a hierarchical prior). In contrast, we use a simple, relatively uninformative, non-hierarchical prior for the centers, *c_i_ ∼* normal(0*, σ*_0_). We set the mean to 0 as the geographic coordinates are centered. We set *σ*_0_ based on the scale of the geographic coordinates, specifically, *σ*_0_ = 3SD(*x*), where SD(*x*) is the standard deviation of the geographic coordinates. We complete the model by specifying priors, which by default are relatively uninformative, for the remaining parameters: *µ_β_ ∼* normal(0*, σ_µ_*), *σ_β_ ∼* gamma(*α* = *α*_0_*, β* = *β*_0_), and *σ_ɛ_ ∼* gamma(*α* = *α*_0_*, β* = *β*_0_). Here, *α*_0_ and *β*_0_ are set by the users (the current defaults are small values, 0.1 and 0.01), and *σ_µ_* is set relative to the scale of the geographic coordinates, *σ_µ_* = 1.5 <u>^4^</u>. As with the genomic cline models, the geographic cline models are fit using HMC with the NUTS algorithm via Stan (Neal *et al*., 2011; Betancourt & Girolami, 2015; Stan Development Team, 2022, 2024). This is done with the est geocl function in bgchm.

### Simulations and results for geogrpahic clines

We analyzed a series of simulated data sets to validate the performance of bgchm’s geographic clines model, with emphasis on (i) the effect of cline variability on inference, and (ii) our ability to estimate cline variability. Similar to our initial tests of the genomic clines models, we simulated data using the geographic clines model as our generative model. This has the key strength of providing known parameter values that can be compared to our estimates. We simulated three levels of cline variability, that is standard deviations in slopes (on the logit-linear scale) among loci (across the genome) of *σ_β_* = 0.1, 0.4 and 0.8. We set the standard deviation for centers (again on the logit-linear scale) to 0.3 and the residual error standard deviation (*σ_ɛ_*) to 0.1. We simulated 100 data sets, each comprising 25 demes and 100 loci, under each of these three levels of cline variability.

We first sampled geographic coordinates for the 25 demes from a standard normal distribution. The resulting coordinates were centered (i.e., forced to have a mean of zero). Next, for each data set, we sampled cline centers and slopes for the 100 loci from normal distributions, *β_i_ ∼* normal(*µ_β_, σ_β_*) and *c_i_ ∼* normal(0, 0.3). Here, *µ_β_* is the mean slope across loci, which we set to 1.75. We then calculated expected logit allele frequencies for each deme and locus as logit(*p_ij_*) = *c_i_* + *β_i_x_j_*; actual logit allele frequencies were sampled from a normal distribution centered on this value but with a standard deviation of *σ_ɛ_*. The resulting allele frequencies (converted to the natural scale), were used as input for bgchm along with the geographic coordinates. We estimated geographic cline parameters with the est geocl function using the default HMC settings of four chains, 2000 total iterations, a 1000 iteration warmup, and a thinning interval of 1.

We found that bgchm produces remarkably accurate and precise estimates of cline parameters, including the standard deviation parameter, which is perhaps unsurprising given the general ease of inference for (hierarchical) linear models (Table S9 and Figure S3). Specifically, the slope standard deviation estimates corresponded closely with the true values of 0.1 (mean across simulations = 0.099), 0.4 (mean = 0.395) and 0.8 (mean = 0.789) (Figure S3B). Moreover, mean absolute errors for slopes and centers for individual loci were uniformly low (means *<* 0.037 and 0.019, respectively) and correlations between true and estimated parameter values were high (mean correlations ranged from 0.925 to 0.998) (Table S9 and Figure S3C-F). Finally, 90% CIs for these parameters contained the true values about the expected proportion of the time, that is about 90% of the time (Table S9)

## Supplemental Tables and Figures

**Table S1:** Mean absolute error (MAE) and 90% credible interval coverage (90% CI cov.) for hybrid index and interpopulation ancestry parameters from bgchm summarized across 50 replicate simulations for each of six sets of conditions. Conditions considered were allele frequency differences (AFDs) of 1, 0.5 or 0.1 with known or uncertain genotypes. Means and standard deviations (in parentheses) across the 50 replicates are given for each metric and set of conditions.

| Conditions | Hybrid index ($H$) | | Interpopulation ancestry ($Q_{10}$) | |
| --- | --- | --- | --- | --- |
|  | MAE | 90% CI cov. | MAE | 90% CI cov. |
| AFD 1.0, known genotypes | 0.002 (0.0002) | 1.000 (0.000) | 0.004 (0.0006) | 1.000 (0.00) |
| AFD 0.5, known genotypes | 0.021 (0.0027) | 0.944 (0.031) | 0.063 (0.0071) | 0.910 (0.039) |
| AFD 0.1, known genotypes | 0.130 (0.0116) | 0.889 (0.040) | 0.161 (0.0161) | 0.976 (0.019) |
| AFD 1.0, uncertain genotypes | 0.004 (0.0005) | 1.000 (0.000) | 0.008 (0.0010) | 1.000 (0.003) |
| AFD 0.5, uncertain genotypes | 0.022 (0.0025) | 0.937 (0.034) | 0.069 (0.0082) | 0.910 (0.041) |
| AFD 0.1, uncertain genotypes | 0.133 (0.0134) | 0.889 (0.051) | 0.166 (0.0196) | 0.977 (0.021) |

**Table S2:**
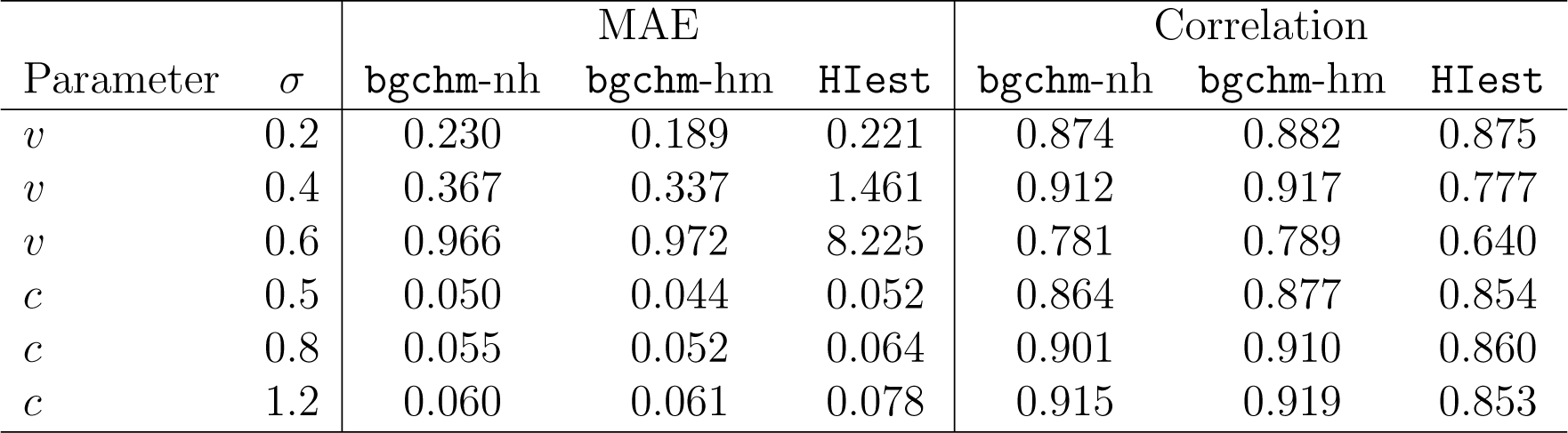
Mean absolute error (MAE) and correlations between true and estimated parameter values for genomic cline slope (*v*) and center (*c*) parameters. Results are shown for different cline standard deviations (*σ_v_* or *σ_c_*) and with inferences based on a non-hierarchical model from bgchm (bgchm-nh), the standard hierarchical model from bgchm (bgchm-hm), or HIest. Means across 50 replicate simulations with fixed allele frequency differences between parents are shown.

| Parameter | $\sigma$ | MAE | | | Correlation | | |
| --- | --- | --- | --- | --- | --- | --- | --- |
|  |  | **bgchm-nh** | **bgchm-hm** | **HIest** | **bgchm-nh** | **bgchm-hm** | **HIest** |
| $v$ | 0.2 | 0.230 | 0.189 | 0.221 | 0.874 | 0.882 | 0.875 |
| $v$ | 0.4 | 0.367 | 0.337 | 1.461 | 0.912 | 0.917 | 0.777 |
| $v$ | 0.6 | 0.966 | 0.972 | 8.225 | 0.781 | 0.789 | 0.640 |
| $c$ | 0.5 | 0.050 | 0.044 | 0.052 | 0.864 | 0.877 | 0.854 |
| $c$ | 0.8 | 0.055 | 0.052 | 0.064 | 0.901 | 0.910 | 0.860 |
| $c$ | 1.2 | 0.060 | 0.061 | 0.078 | 0.915 | 0.919 | 0.853 |

**Table S3:** Proportion of loci with 90% credible intervals (CIs) overlapping the true value or 0 (the null expectation) for genomic cline slope (*v*) and center (*c*) parameters. Results are shown for different cline standard deviations (*σ_v_* or *σ_c_*) and with inferences based on a non-hierarchical model from bgchm (bgchm-nh) or the standard hierarchical model from bgchm (bgchm-hm). Means across 50 replicate simulations with fixed allele frequency differences between parents are shown.

| Parameter | $\sigma$ | CI overlap true value | | CI overlap 0 | |
| --- | --- | --- | --- | --- | --- |
|  |  | **bgchm-nh** | **bgchm-hm** | **bgchm-nh** | **bgchm-hm** |
| $v$ | 0.2 | 0.89 | 0.89 | 0.53 | 0.58 |
| $v$ | 0.4 | 0.88 | 0.87 | 0.30 | 0.31 |
| $v$ | 0.6 | 0.81 | 0.78 | 0.21 | 0.22 |
| $c$ | 0.5 | 0.90 | 0.87 | 0.55 | 0.61 |
| $c$ | 0.8 | 0.89 | 0.87 | 0.43 | 0.46 |
| $c$ | 1.2 | 0.90 | 0.84 | 0.36 | 0.38 |

**Table S4:** Mean absolute error (MAE), correlations between true and estimated parameter values, and proportion of loci with 90% credible intervals (CIs) overlapping the true value for genomic cline slope (*v*) and center (*c*) parameters. Conditions considered were minimum allele frequency differences (AFDs) of 1, 0.5 or 0.1 with known or uncertain genotypes. Means across 50 replicates for each set of simulation conditions are shown.

| Conditions | MAE |  | Correlation |  | CI overlap 0 |  |
| --- | --- | --- | --- | --- | --- | --- |
| | $v$ | $c$ | $v$ | $c$ | $v$ | $c$ |
| AFD 1.0, known genotypes | 0.24 | 0.05 | 0.93 | 0.91 | 0.88 | 0.87 |
| AFD 0.5, known genotypes | 0.43 | 0.07 | 0.77 | 0.85 | 0.87 | 0.86 |
| AFD 0.1, known genotypes | 0.68 | 0.09 | 0.60 | 0.71 | 0.87 | 0.81 |
| AFD 1.0, uncertain genotypes | 0.26 | 0.05 | 0.92 | 0.90 | 0.88 | 0.88 |
| AFD 0.5, uncertain genotypes | 0.45 | 0.07 | 0.76 | 0.84 | 0.87 | 0.85 |
| AFD 0.1, uncertain genotypes | 0.71 | 0.09 | 0.58 | 0.69 | 0.87 | 0.80 |

**Table S5:** Numbers of loci with credible deviations from genome-average introgression based on 90% equal-tail probability intervals. Results are summarized for ten replicates each of neutral, oligogenic (two underdominant loci), and weak and strong polygenic (50 underdominant loci) selection and for cline gradient (*v*) and center (*c*). Means and standard deviation s (SDs) across the ten replicates are reported; numbers are out of 251 loci.

| Conditions | gradient ($v$) | | center ($c$) | |
| --- | --- | --- | --- | --- |
|  | mean | SD | mean | SD |
| Neutral | 74.2 | 4.4 | 184.4 | 3.9 |
| Oligogenic | 68.5 | 5.6 | 175.9 | 8.7 |
| Weak polygenic | 74.6 | 5.2 | 184.9 | 5.7 |
| Strong polygenic | 68.7 | 6.5 | 168.2 | 7.4 |

**Table S6:** Correlation between point estimates of log cline gradient parameters (*v*) and the distance in centimorgans to the nearest underdominant locus. Results are summarized for ten replicates each of neutral, oligogenic (two underdominant loci), and weak and strong polygenic (50 underdominant loci) selection. For neutral simulations, there were not underdominant loci, and thus we used the underdominant locus positions from the polygenic simulations (this is done to verify observed patterns with selection are not artifactual). Cases where the reported correlation is significantly different from 0 (*P <* 0.05) are in bold.

| Replicate | Neutral | Oligogenic | Weak<br>polygenic | Strong<br>polygenic |
| --- | --- | --- | --- | --- |
| 1 | 0.05 | **-0.36** | -0.07 | **-0.16** |
| 2 | 0.04 | **-0.47** | -0.11 | **-0.19** |
| 3 | 0.04 | **-0.48** | -0.08 | **-0.11** |
| 4 | 0.00 | **-0.37** | -0.08 | **-0.14** |
| 5 | 0.06 | **-0.48** | -0.07 | **-0.20** |
| 6 | -0.02 | **-0.48** | -0.10 | **-0.16** |
| 7 | 0.05 | **-0.42** | -0.06 | **-0.13** |
| 8 | 0.01 | **-0.49** | -0.09 | **-0.18** |
| 9 | 0.03 | **-0.45** | -0.04 | **-0.13** |
| 10 | 0.03 | **-0.34** | -0.08 | **-0.23** |

**Table S7:** Correlation between point estimates of the absolute value of logit cline gradient centers (*c*) and the distance in centimorgans to the nearest underdominant locus. Results are summarized for ten replicates each of neutral, oligogenic (two underdominant loci), and weak and strong polygenic (50 underdominant loci) selection. For neutral simulations, there were not underdominant loci, and thus we used the underdominant locus positions from the polygenic simulations (this is done to verify observed patterns with selection are not artifactual). Cases where the reported correlation is significantly different from 0 (*P <* 0.05) are in bold.

| Replicate | Neutral | Oligogenic | Weak<br>polygenic | Strong<br>polygenic |
| --- | --- | --- | --- | --- |
| 1 | 0.03 | **0.20** | 0.04 | **0.18** |
| 2 | 0.01 | **0.28** | 0.07 | **0.17** |
| 3 | -0.01 | **0.25** | 0.05 | 0.08 |
| 4 | -0.02 | **0.15** | 0.07 | **0.13** |
| 5 | 0.01 | **0.25** | 0.05 | **0.15** |
| 6 | 0.02 | **0.29** | 0.10 | **0.13** |
| 7 | 0.03 | **0.29** | 0.05 | 0.11 |
| 8 | 0.01 | **0.19** | 0.03 | **0.13** |
| 9 | -0.01 | **0.17** | 0.09 | **0.13** |
| 10 | 0.02 | **0.28** | 0.03 | **0.17** |

**Table S8:** Summary of evidence for an excess of loci deviating from genome-wide introgression on the Z chromosome. Deviations considered include *v >* 1 (steeper clines), *v <* 1 (shallower clines), center *>* 0.5 (excess Jackson Hole *Lycaeides* ancestry) and center *<* 0.5 (excess *L. melissa* ancestry). Classifications are based on the 90% credible intervals for each parameter excluding the relevant null value (1 or 0.5) in the specified direction. The total number of such loci, the number on the Z, and the number on the Z expected by chance given the number of autosomal and Z SNPs analyzed (based on the mean of 1000 randomizations) are given, along with the associated *P* -value for the number on the Z exceeding the null expectations from the 1000 randomizations.

| Condition | Total | Z observed | Z expected | $P$-value |
| --- | --- | --- | --- | --- |
| $v > 1$ | 40 | 39 | 13.7 | 0.001 |
| $v < 1$ | 60 | 12 | 40.5 | 0.999 |
| center $> 0.5$ | 48 | 23 | 16.4 | 0.022 |
| center $< 0.5$ | 70 | 37 | 23.8 | 0.001 |

**Table S9:** Mean absolute error (MAE), correlations between true and estimated parameter values, and proportion of loci with 90% credible intervals (CIs) overlapping the true value for geographic cline slope (*β*) and center (*c*) parameters. Results are shown for slope standard deviations of 0.1, 0.4 and 0.8 Means across 100 replicates for each set of simulation conditions are shown.

| $\sigma$ | MAE | | Correlation | | CI overlap 0 | |
| --- | --- | --- | --- | --- | --- | --- |
| | $\beta$ | $c$ | $\beta$ | $c$ | $\beta$ | $c$ |
| 0.1 | 0.031 | 0.017 | 0.925 | 0.997 | 0.889 | 0.904 |
| 0.4 | 0.034 | 0.018 | 0.994 | 0.997 | 0.893 | 0.904 |
| 0.8 | 0.036 | 0.018 | 0.998 | 0.997 | 0.891 | 0.901 |

**Figure S1:**
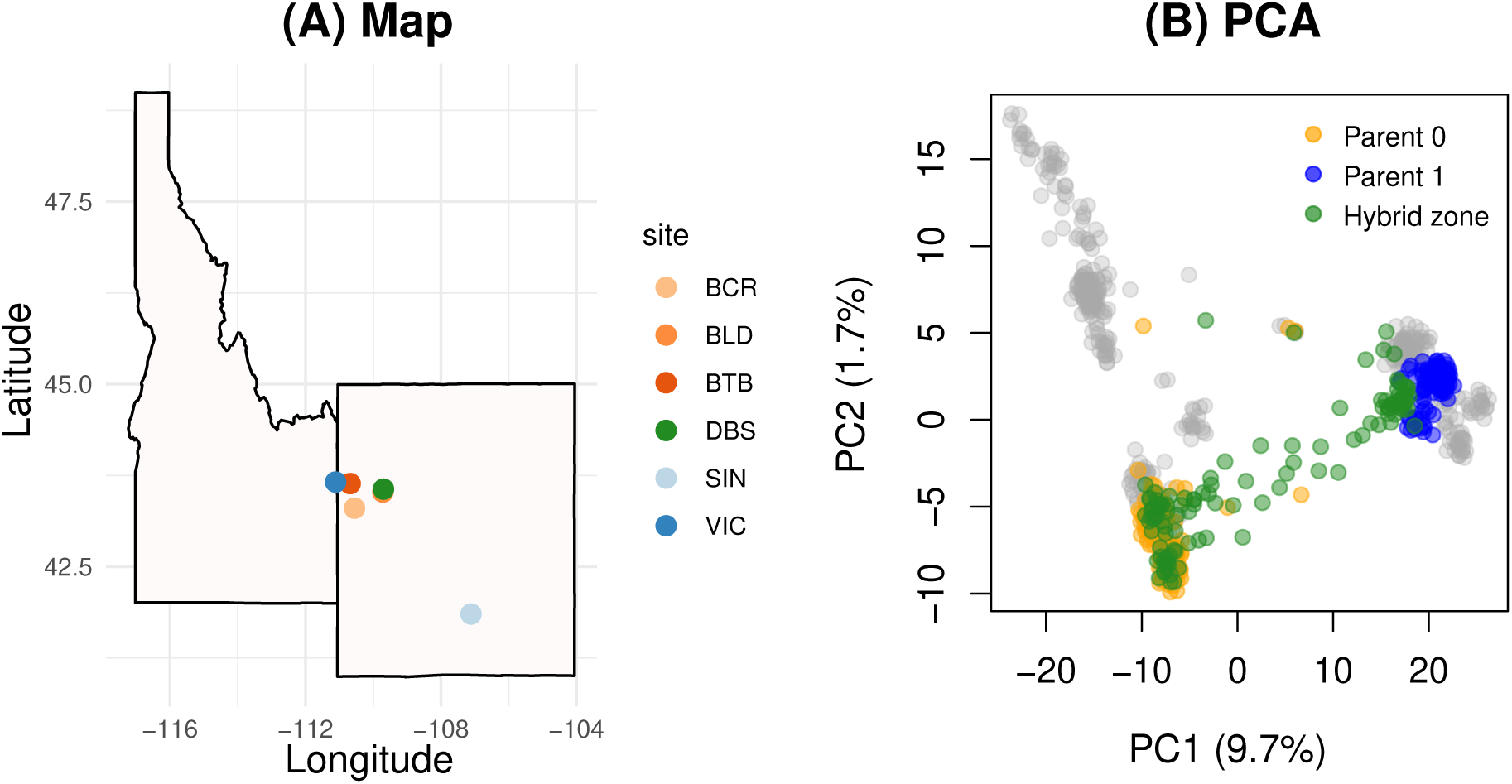
Map and summary of genetic variation for the *Lycaedies* hybrid zone. The map shows outlines for the US states of Idaho and Wyoming along with points for the focal *Lycaeides* populations in this paper (A). This includes the two *L. melissa* populations (SIN and VIC), the three Jackson Hole *Lycaeides* populations (BCR, BLD and BTB), and the hybrid zone population (DBS). Panel (B) summarizes patterns of genetic variation with a principal components analysis (PCA). The first two PC axes are shown. Each point denotes an individual butterfly. Orange and blue points correspond with Jackson Hole *Lycaeides* and *L. melissa* reference source populations, respectively. Green points denote individuals form the Dubois hybrid zone. Gray dots indicate butterflies not included in the hybrid zone analysis, including non-admixed *L. idas*.

**Figure S2:**
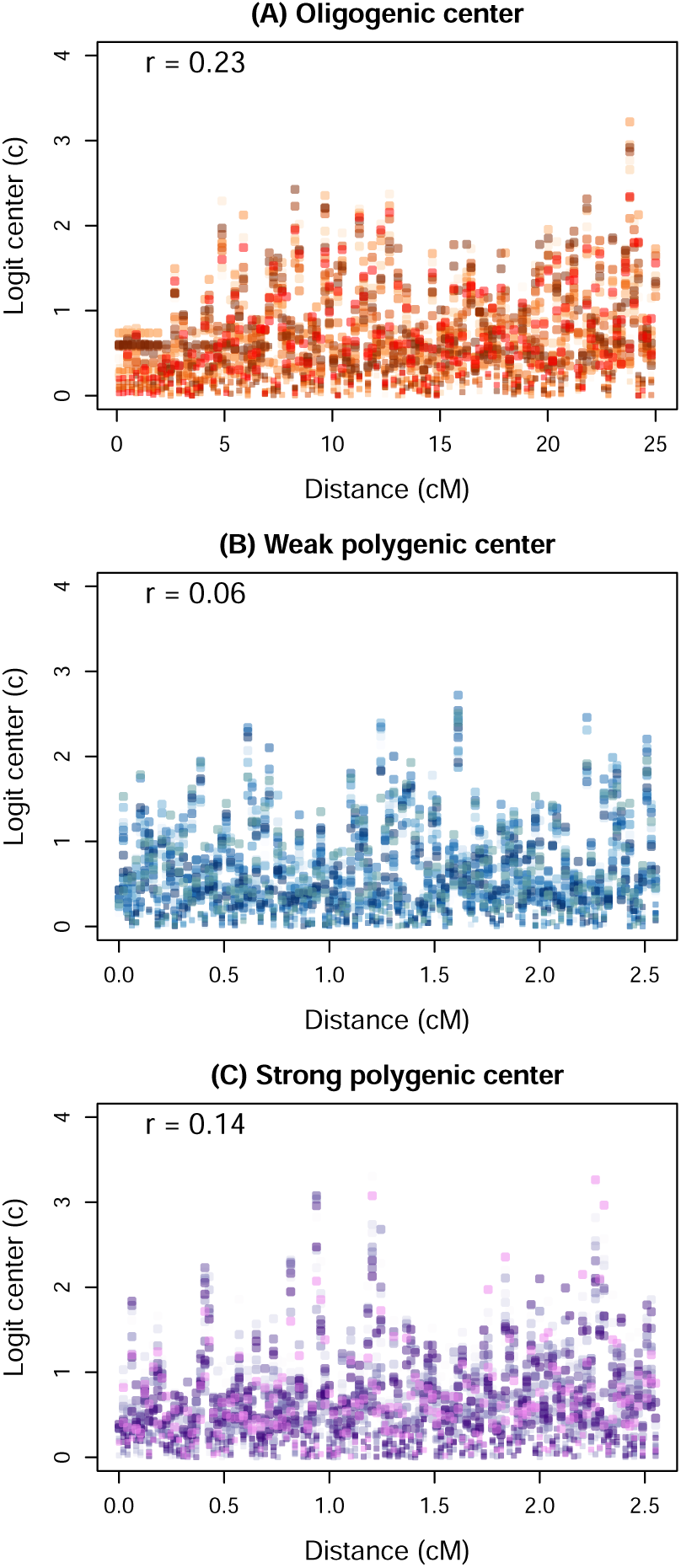
Summary of genomic cline center estimates for hybrid zone simulations with alternative genetic architectures for hybrid fitness. Results are shown for oligenic selection (A), weak polygenic selection (B) and strong polygenic selections (C) (see main text for details). Plots show the relationships between the distance (in cM) a marker locus is from a selected locus and the logit of the cline center (*c*). This is only shown for the three sets of conditions with selection. Points are colored to indicate distinct replicate simulations and the Pearson correlation between distance and logit(*c*) is reported.

**Figure S3:**
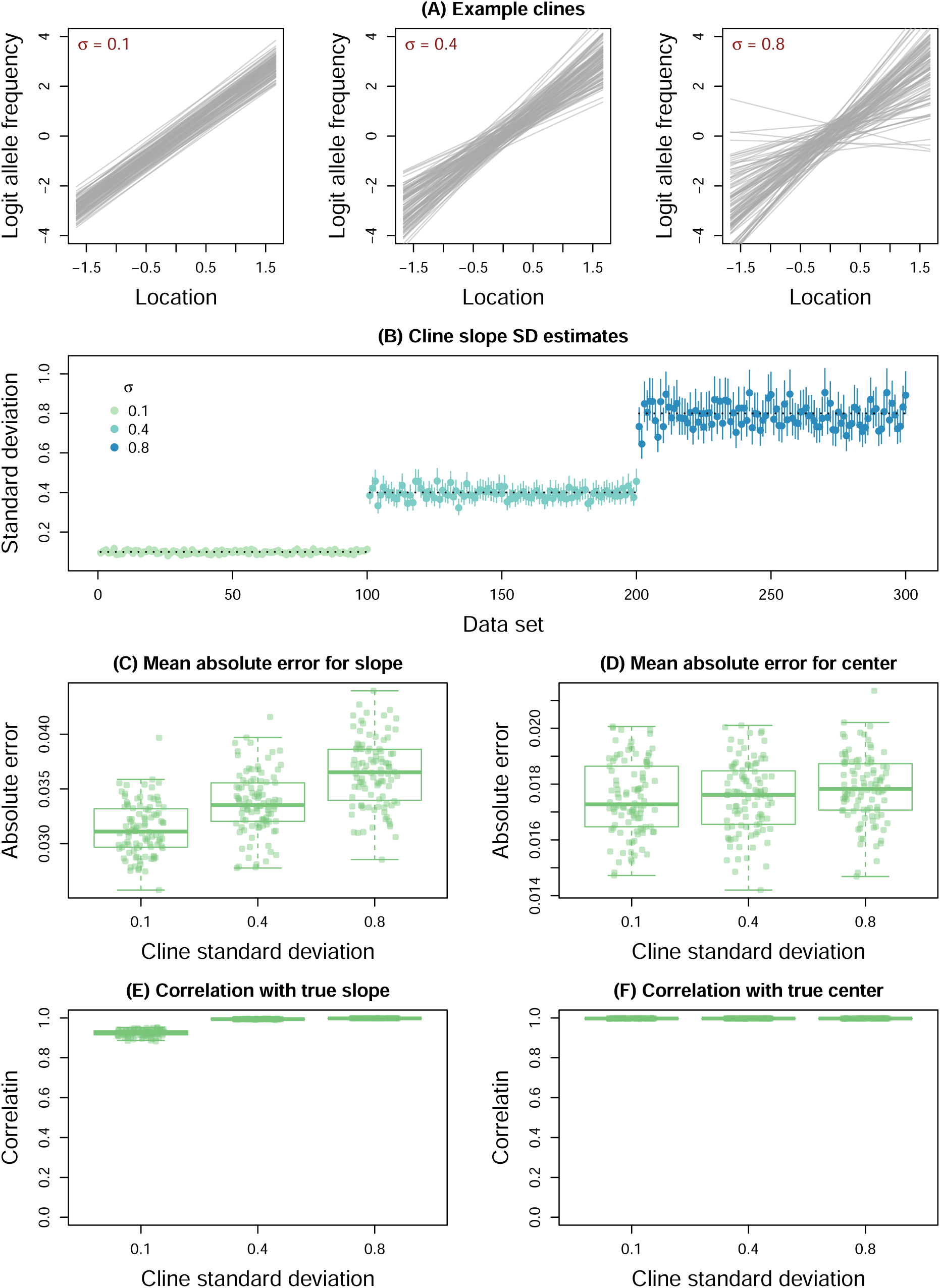
Summary of geographic cline variability and the effect of such variability on cline inference. Panel (A) shows geographic cline estimates with low (*σ* = 0.1), moderate (*σ* = 0.4), and high (*σ* = 0.8) variability in slopes on the logit scale. Each gray line is the cline for a locus and gives the expected logit allele frequency as a function of geographic location. Estimates of cline standard deviations for slopes are shown in panels (B). Here, point estimates and 90% credible intervals (CIs) are depicted with points and vertical lines. Horizontal dotted lines give the true value used for each simulation. Performance, in terms of estimating cline slopes (*β*) and centers, is summarized based on mean absolute error in panels (C) and (D) and in terms of the correlation between true and estimated parameter values in panels (E) and (F). Errors and Pearson correlations were computed based on parameter point estimates (posterior medians) and are summarized across replicate simulations with boxplots. Boxes indicate the median and 1st and 3rd quartiles of the distribution across replicate simulations, with whiskers extending up to 1.5*×* the interquartile range. The overlain points show metrics for individual replicates.

